# Weaning imprints a hybrid cytotoxic-regulatory identity on intestinal CD4^+^ T lymphocytes

**DOI:** 10.64898/2026.10.26.762022

**Authors:** Marina Schernthanner, Rohit R. Gokhale, Alexandra Thoms, Gabriella Lima dos Reis, Marwa Saad, Mackenzie M. Clark, Yue S. Yin, Izabela Mamede, Bernardo S. Reis, Tiago B. R. Castro, Roham Parsa, Elaine Fuchs, Daniel Mucida, Angelina M. Bilate

## Abstract

The intestinal immune system must adapt as the neonatal gut shifts from milk to solid food and becomes densely colonized. How the T cell compartment is established under these conditions remains unclear. Using fate-mapping to timestamp naïve T cells, with imaging, transcriptomic, and repertoire analyses, we tracked them from suckling to adulthood. Intestinal T cell entry peaked at weaning, coinciding with relocation from crypts to villi guided by the tissue microenvironment and, partly, the microbiota. T cells timestamped at suckling persisted into adulthood and, at weaning, adopted a transient cytotoxic-regulatory state in the lamina propria, with granzyme B expression microbiota-dependent only in the ileum. Clonal expansions at weaning were restricted to the small intestinal lamina propria and absent in germ-free mice. Most expanded clones recognized early-life commensals. Weaning is a physiological window of transient inflammation imprinting a hybrid CD4⁺ T cell compartment, suggesting that early-life commensals modulate T cell selection.

## Introduction

The gut mucosa hosts one of the most active sites of adaptive immunity at steady state, continuously interacting with a diverse, shifting mixture of dietary and microbial antigens. Adaptive immunity deficiency impairs intestinal homeostasis, even in the absence of infection^1–3^, suggesting that this engagement serves organismal physiology, not only host defense.

At birth, mammals transition from a largely sterile uterine environment to one in which microbes immediately colonize barrier tissues. During suckling, microbes that thrive on an exclusive milk diet predominate in the intestinal lumen. During this period, CD4^+^ T cells enter the gut, particularly in lymphoid structures such as Peyer’s patches^4^. At weaning, the introduction of solid food coincides with increased microbial diversity and density, accompanied by a microbiota-dependent immunological process termed the “weaning reaction”^5^, which exposes the developing intestinal immune system to a suite of new antigens. However, how T lymphocytes seed the intestine and adapt to lifelong microbial colonization remains poorly understood.

Addressing this question requires an *in vivo* system that is physiologically colonized and allows the tracking of defined antigen specificities. However, most mouse models achieve one at the expense of the other. Monoclonal TCR models resolve single antigen specificity and have shown that commensal-specific T cells regulate intestinal immunity^6–10^, but they do not capture the polyclonal response to the endogenous microbiota. Defined microbial consortia, such as Oligo-MM^12^ ^11^ and hCom^10,12^ have been instrumental in dissecting the contribution of individual taxa, but they do not fully recapitulate the natural complexity of the microbiome. For example, mice colonized with Oligo-MM¹² or hCom show intestinal T cell responses that differ, to some extent, from those of specific pathogen-free (SPF) mice^10,13^, likely because of lower microbial diversity or poor adaptation of the bacterial strains to the mouse gut. In contrast, wild or pet store mice, whose microbiota approximates natural complexity but also includes infectious pathogens, develop stronger T cell effector responses than SPF mice^14,15^. These findings suggest that microbial diversity, not only colonization by itself, calibrates T cell differentiation in the gut, a graded relationship that simplified systems only partially capture.

To pair physiological colonization with clonal resolution, we used inducible fate-mapping to “timestamp” naïve T cells in SPF mice at defined postnatal stages and followed them from suckling to adulthood by imaging, transcriptomics, and TCR sequencing. T cell entry peaks at weaning, a window in which T cells relocate from crypts to villi. In the small intestinal lamina propria, weaning also coincides with microbiota-dependent clonal expansion and a transient hybrid cytotoxic-regulatory state. At the clonal level, we linked recognition of early-life commensals to this state in expanded clones.

## Results

### Intestinal T lymphocytes redistribute from crypt to villus during postnatal development

Whole-tissue analyses have previously described an early wave of adaptive lymphocytes that steadily increased after birth^4^. To first resolve this by compartment, we profiled the mononuclear cells of the small intestine by flow cytometry across postnatal development, distinguishing the intestinal epithelium (IE) and lamina propria (LP) compartments. Whereas innate populations, especially macrophages and innate lymphoid cells (ILCs), dominated during suckling (<P14) in the LP, a wave of αβT cells surpassed those at one week after birth. This agrees with reports that T lymphocytes outcompete ILCs and NKT cells in the gut^16,17^. After a sudden drop in the relative frequency of αβT cells around the two-week mark, they rose steadily after weaning (Figures 1A and S1A). These dynamics of αβT cells in the LP were largely mirrored in the IE with a delay, and were accompanied by γδT cells peaking after birth and again at weaning. Overall, the innate populations remained stable whereas the adaptive subsets fluctuated with age.

**Figure 1.**
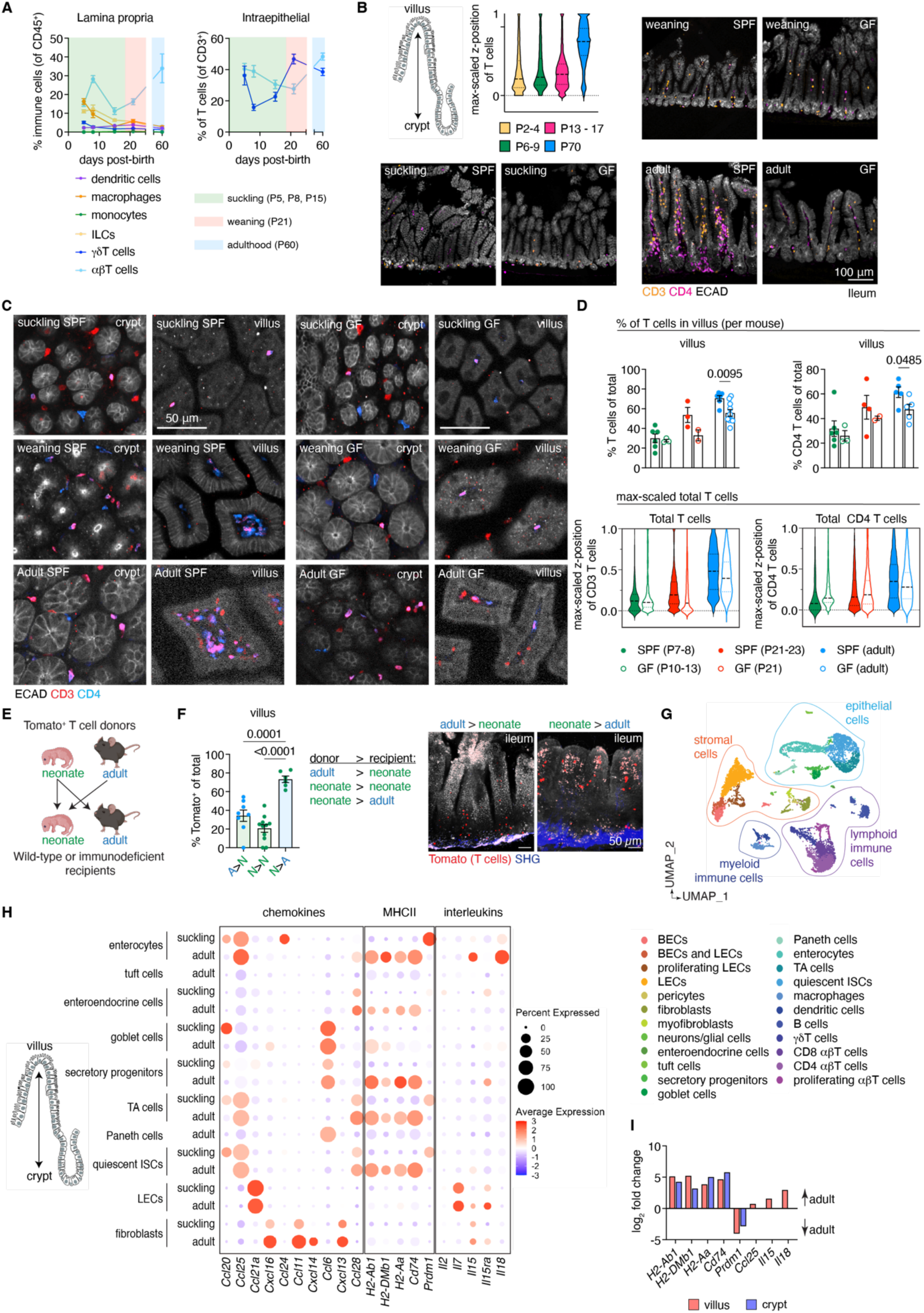
Intestinal T lymphocytes redistribute from crypt to villus during postnatal development. **(A)** Profiling of innate and adaptive immune subsets in the small intestinal lamina propria and intraepithelial compartment of specific-pathogen-free (SPF) mice of the indicated ages (n = 3-8 mice/age). **(B)** Schematic of the crypt–villus axis (left). Max-scaled z-position of individual ileal T cells along the axis quantified from whole-mount samples. Sagittal ileal sections from SPF and germ-free (GF) mice at suckling (P7), weaning (P18) and adult (8 weeks) stages, CD3ε^+^ (orange) and CD4^+^ (pink) T cells. **(C)** Representative ileal whole-mount z-slice images at the crypt (left) and villus (right) level from suckling, weaning and adult SPF and GF mice. (**D**) Quantification of location of CD3^+^ (left) and CD4^+^ (right) T cells in (C). Top: fraction of T cells in the villus per mouse. Bottom: max-scaled z-position of individual T cells along the crypt-to-villus axis, pooled from all mice per age group (n=2-10 mice/age). **(E-F)** Heterochronic adoptive transfer. Total CD4^+^ cells were isolated from the mesenteric lymph node and spleen of Tomato^+^ neonatal (P7) or adult donors and transferred into neonatal or adult *Rag1*^-/-^, *Rag2*^−/−^, or CD45.1 recipients in separate experiments. **(E)** Quantification of Tomato^+^ T cells in the villus 1 week after transfer **(F)**. Ileum of *Rag1*^−/−^ recipients. Tomato^+^ T cells (red), second harmonic generation (blue), and autofluorescence (white); n = 6-11 mice per condition. (**G**) UMAP of 10X scRNA-seq from integrated data of the small intestine of neonatal (P6-7) and adult mice (Adult set from GSE190037). BECs, blood endothelial cells; LECs, lymphatic endothelial cells; ISCs, intestinal stem cells; TA, transit-amplifying cells. (**H**) Dot plot of selected chemokines, MHC-II, and interleukin genes in non-immune cells of suckling and adult mouse small intestine, ordered by spatial position along the crypt–villus axis. Tuft and Paneth cells were only recovered in sufficient numbers in adult samples. (**I**) Log2-fold change of differentially expressed genes (DEG) in villus and crypt epithelial cells, adult vs. suckling mice. Genes without a bar did not meet adjusted *p* < 0.05 and |log2FC| > 0.585. Each symbol represents a mouse; mean ± SEM. Unpaired two-tailed t-test, SPF vs. GF mice per timepoint (D). Data pooled from 9 independent experiments (E-F); one-way ANOVA with Tukey’s test (F).

We complemented flow cytometry with imaging to position immune cells within the developing gut, in the presence or absence of microbial colonization. Multiplex imaging by iterative bleaching extends multiplexity (IBEX)^18^ immunostaining of the terminal ileum showed myeloid cells evenly dispersed throughout the tissue at suckling, weaning, and adult timepoints (Figure S1B), consistent with their stable frequencies detected by flow cytometry. Although a large proportion of αβT cells occupy Peyer’s patches in the neonatal intestine^4^, we readily detected T cells in the non-lymphoid tissue of suckling mice. T cells preferentially occupied intestinal crypts before gradually redistributing along the villi after weaning, a pattern that was less evident in germ-free (GF) mice (Figures 1B and S1C-D). To gain a more granular view of these dynamics, we employed 3D-immunofluorescence imaging of cleared whole-mount ileal samples from SPF and GF mice. In both SPF and GF suckling mice, T cells localized to the crypt, but migration to the villus at weaning and beyond was blunted in GF mice (Figure 1C-D; Video S1-2), suggesting that the microbiota only partially drives T cell distribution in the tissue.

Next, we used heterochronic adoptive transfer of neonatal or adult T cells (Figure 1E) to test whether T cell localization was dictated by T cell-intrinsic programs or the tissue microenvironment. Regardless of donor age, T cells were localized around the crypts in neonatal recipients and along the villi in adult recipients (Figure 1F), indicating that the tissue microenvironment is the major driver of T cell distribution in the intestine. To identify the molecular cues guiding T cell positioning, we profiled the neonatal (P6) small intestine using single-cell RNA sequencing (scRNA-seq), enriching for T cells and crypt epithelial cells to offset their underrepresentation relative to the villus epithelium (Figures 1G and S1E). In line with our immune profiling studies, CD4^+^ T cells dominated the T cell compartment and exhibited a naïve phenotype (Figure S1F). Integrating this dataset with a published scRNA-seq dataset of the adult mouse small intestine^19^ allowed us to map age-related changes in chemokines, antigen presentation machinery, and cytokines across epithelial and stromal populations ordered along the crypt-villus axis (Figures 1G and S1E).

Transcriptional programs were largely conserved across cell types and age, but the expression of antimicrobials (C-type lectins and defensins), recruitment signals (*Ccl25*), interleukins (*Il15*, *Il18*), and genes encoding major histocompatibility complex (MHC) II molecules (*H2-Ab1*, *H2-DMb1*, *H2-Aa*, *Cd74*) increased with age (Figures 1H-I and S1G). *Ccl25*, the dominant gut-homing chemokine, was expressed by transit-amplifying (TA) cells and intestinal stem cells (ISCs) of the crypt at comparable levels between suckling and adult mice (Figures 1H and S1H) but increased in adult villus enterocytes with age. The expression of *Il18* and the T cell survival factor *Il15* was also elevated in adult villus epithelial cells (Figure 1I). The genes of the MHC-II pathway, as well as MHC-II expression by IE cells, gradually increased in the crypt and villus epithelium after weaning in a microbiota-dependent manner (Figures 1H-I and S1I-L). Conversely, *Prdm1* (Blimp-1), which has been implicated in the repression of MHC-II in immune cells^20,21^ and in regulating epithelial maturation during the suckling-to-weaning transition^22^, was elevated in suckling mice (Figure 1H-I). Overall, our data show that T cells occupy discrete niches during postnatal development in mice. We propose that with age, epithelial maturation programs transform villus epithelial cells into a rich source of homing (*Ccl25*) and survival (*Il15*) factors for T cells, while MHC-II on epithelial cells, induced by microbial colonization, may engage and maintain CD4^+^ T cells in the intestinal mucosa. In the absence of such programs, as is the case of the intestine of neonates and germ-free mice, T cells appear to linger around epithelial crypts by default.

### T cell recruitment to the small intestine peaks during weaning

Our findings above suggested that T cells can enter the suckling gut while still expressing CD62L. Therefore, we profiled intestinal αβT cells from the lamina propria and epithelium of small (SI) and large intestine (LI) during suckling, weaning, and adulthood. CD4^+^ T cells dominated the TCRαβ^+^ compartment during suckling (P7, P14) relative to CD8αβ^+^ cells in both LP and IE of SI and LI; this ratio shifted in the epithelium at weaning and equalized by adulthood (Figures 2A and S2A-B). Most intestinal T cells expressed CD62L during suckling (P7, P14), but lost it by weaning (P17, P22) (Figure 2B). These CD62L^+^ T cells in the neonatal gut were not blood contaminants as they remained largely unlabeled in mice injected intravenously with fluorescently labeled anti-CD45 (Figure S2C). Although intestinal T cells expressed α4β7 at similar levels throughout postnatal development (Figure S2D), the density of its ligand mucosal vascular addressin cell adhesion molecule 1 (MAdCAM-1) on blood endothelial cells peaked in the suckling intestine (Figure 2C). Increased MAdCAM-1 density in the neonatal intestine may facilitate the entry of CD62L^+^ T cells near the crypt during the suckling period. In summary, naïve CD4^+^ and CD8^+^ T cells seeded the small and large intestine in the first weeks of life, when MAdCAM-1 density and CD62L expression peaked, whereas weaning coincided with a shift to CD62L^-/low^ T cells, leaving α4β7-MAdCAM-1 binding as the major route for continued T cell entry in adulthood.

**Figure 2.**
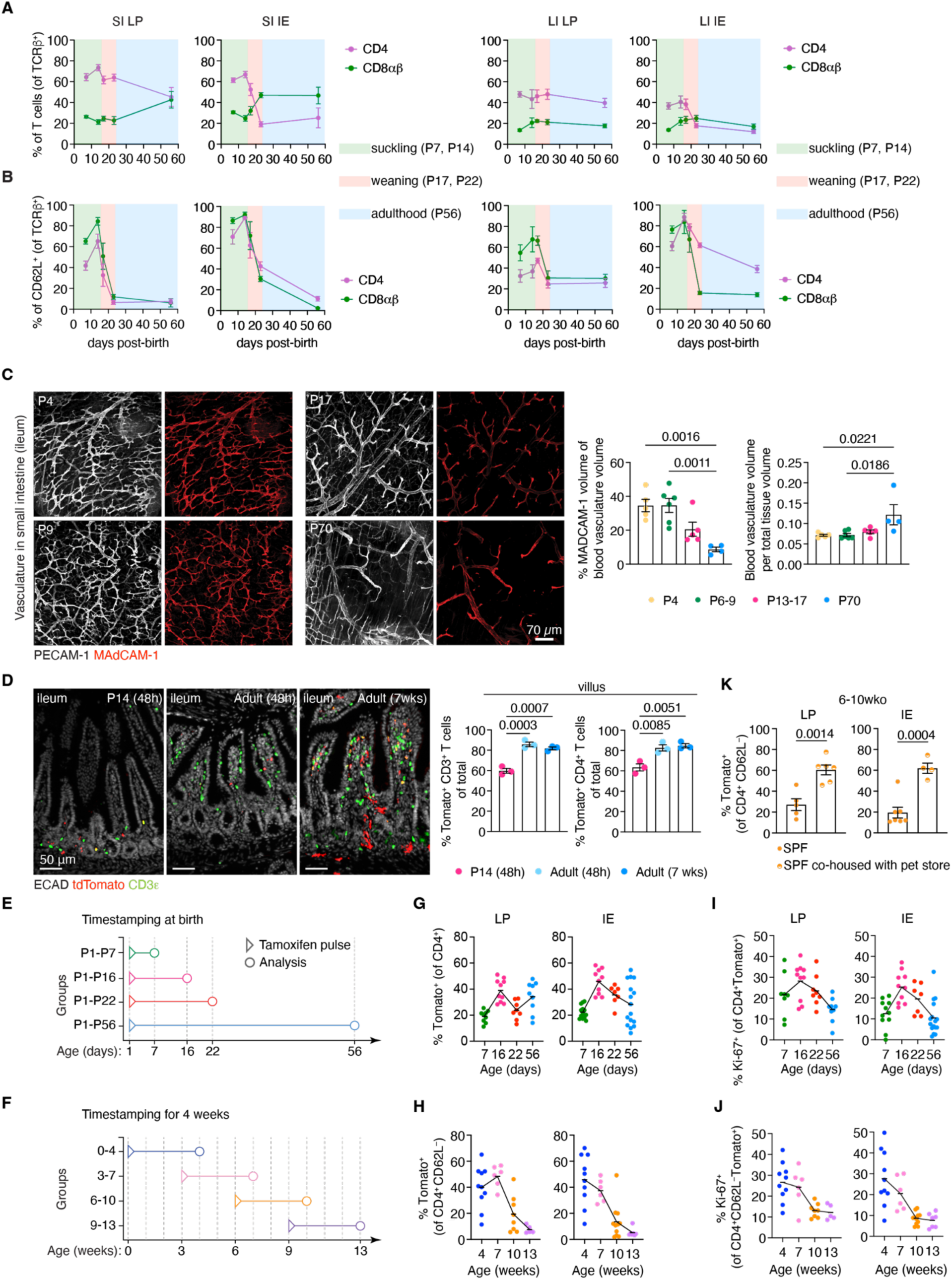
T cell recruitment to the small intestine peaks during weaning. **(A-B)** Proportion of the indicated αβT cells (A) or CD62L^+^ naïve αβT cells (B) across postnatal ages in SPF mice. SI, small intestine; IE, intraepithelial lymphocytes; LP, lamina propria lymphocytes; LI, large intestine. (n=3-9 mice/age). **(C)** Representative max-projected z-stack images of the blood vasculature (PECAM-1, white; MAdCAM-1, red) below the crypts in ileal whole-mount samples of SPF mice. Quantification of MAdCAM-1^+^ structures relative to vascular volume and vascular volume per tissue volume (right graphs). (n=4-6 mice/age). **(D)** Sagittal sections (ileum) of P14 and adult SPF Sell-Tomato mice 48h or 7 weeks after tamoxifen, as indicated in the images (left). Percentage of all Tomato^+^ or Tomato^+^ CD4^+^ T cells in the villus (right). (n=3 mice/age). **(E)** Schematic of timestamping windows in Sell-Tomato mice for graphs in G, I. **(F)** Schematic of timestamping windows in Sell-Tomato mice for graphs in H, J. **(G-H)** Frequency of Tomato^+^ cells in the LP (left) and IE (right). (G) out of CD4^+^ T cells. (H) out of CD4^+^ CD62L^-^ T cells. (n=4-16 mice/age). **(I-J)** Frequency of Ki-67^+^ cells in the LP (left) and IE (right). (I) out of CD4^+^ Tomato^+^ T cells. (H) out of CD4^+^ CD62L^-^ Tomato^+^ T cells. (n=4-16 mice/age). **(K)** Percentage of timestamped CD4^+^ T cells in adult SPF mice co-housed or not with pet-store mice (6-10 wko window). (n=4-7 mice/age). (A, B) Curves show mean ± SEM of all mice analyzed. Bar plots display the mean ± SEM; each symbol represents one mouse. *P* values were calculated by unpaired Student’s t-test for groups of 2 and one-way ANOVA with Tukey’s multiple comparisons for more than 2 groups.

To determine whether early-life T cells persist in the intestine and which fates they acquire, we used inducible fate-mapping to timestamp naïve T cells throughout postnatal development. Tamoxifen treatment of *Sell*^CreERT2^ X *Rosa26*^CAG-LSL-tdTomato^ mice permanently labels naïve T cells expressing L-selectin (*Sell*, CD62L^+^) as Tomato^+^ ^23,24^. Tomato labeling, assessed 24 h after the last tamoxifen dose, was below 10% in thymocytes and splenocytes of neonates and adults and highest in gut-draining mesenteric lymph nodes (mLN) (Figure S2E), where nearly all labeled cells expressed CD62L, confirming specificity (Figure S2F). In the small intestine, labeling at 24 h was modest but higher in neonates than in adults (Figure S2E-F), consistent with CD62L expression on intestinal T cells of neonates (Figure 2B). At 96 h post-tamoxifen, however, intestinal Tomato^+^ CD4^+^ T cells were significantly more frequent in neonates than in adults (Figure S2G). Because the tissue-specific differences exceeded the modest labeling already present at 24h, labeling efficiency alone cannot account for this gap. In the intestine, more Tomato^+^ cells expressed CD62L in the IE than in the LP (Figure S2H), and intravenous anti-CD45 labeling confirmed that timestamped T cells were not blood contaminants (Figure S2I). Confocal imaging localized timestamped T cells along the crypt-to-villus axis during suckling and adulthood. Whereas in P14 mice, Tomato^+^ T cells detected 48h after tamoxifen pulse localized predominantly to the crypt, in adult mice, Tomato^+^ T cells localized to the villus (Figure 2D). When tamoxifen was administered at P10 and mice were analyzed 7 weeks later, Tomato^+^ T cells accumulated in the villus. Thus, the tissue microenvironment appears to dictate where naïve T cells reside and to support the long-term maintenance of early-life arrivals in the villus. These data validated the applicability and specificity of our timestamping approach.

To determine whether the accumulation of timestamped T cells after the 7-week chase reflected staggered waves of incoming T cells or local expansion, we devised two complementary tamoxifen pulse-chase schemes (Figure 2E-F). In the first scheme, the pulse was given at birth (P1) and the intestinal T cell pool was analyzed at suckling (P7 or P16), weaning (P22) and adulthood (P56) (Figure 2E), quantifying the contribution of neonatal T cells to the tissue at defined later stages. In the second scheme, we fixed the pulse-to-analysis interval at four weeks, timestamping naïve T cells at P1 (week 0-4), weaning (week 3-7), or adulthood (week 6-10 and 9-13), (Figure 2F) to quantify the age-dependent entry rate of intestinal T cells. Naïve T cells timestamped at birth contributed substantially to the LP and IE during the suckling-to-weaning transition and persisted for at least 8 weeks in both compartments (Figure 2G), indicating that the intestinal T cell compartment is not entirely replaced by latecomers. In the 4-week scheme, entry peaked in the window spanning weaning (3-7wk) and declined progressively thereafter in both LP and IE (Figure 2H).

In the P1-pulse scheme, the proportion of proliferating Ki-67^+^ T cells among timestamped Tomato^+^cells was smaller than in Tomato^-^ cells, consistent with the Tomato⁺ compartment representing an increasingly resident cohort, while Tomato^-^ cells likely included recent arrivals from the mLN (Figure S2J). This difference was absent in the 4-week-window scheme, where Tomato^+^ cells were a recent cohort at every age (Figure S2K). Overall, proliferation among timestamped T cells was highest in windows spanning birth to weaning and declined thereafter (Figure 2I-J). These trends may reflect homeostatic proliferation driven by neonatal lymphopenia^25^ and/or a burst of proliferation driven by weaning-associated changes in diet and microbial antigen diversity that likely stabilizes in adults. The age-dependent decline in T cell entry did not reflect intrinsic inability of adult T cells to migrate to and persist in the intestine, as co-housing adult SPF mice with pet store mice^14^ significantly increased the fraction of timestamped intestinal T cells (Figure 2K). Hence, T cell entry and proliferation in the small intestine peaked at weaning and declined progressively with age but remained responsive to new microbial exposure in adulthood.

### Weaning establishes a hybrid cytotoxic-regulatory program on lamina propria T lymphocytes

To determine how timestamped T cells transcriptionally adapt to the developing intestine, we sorted CD4^+^CD62L^-/low^ Tomato^+^ T cells from mLN, SI LP and IE from four cohorts used for fate-mapping shown in Figure 2: P1-P16 (suckling, birth-pulse scheme), P1-P22 (weaning, birth-pulse scheme), 3-7wko (weaning-adult, 4-week-window scheme), and 6-10wko (adult-adult, 4-week-window scheme) and performed single-cell RNA-and TCR-sequencing (10X Genomics) (Figure 3A; Table S1). P1-P22 and 3-7wko independently captured the weaning transition using different pulse schemes, and 6-10wko provided an adult steady-state comparison. CD1d tetramer staining was used to exclude NKT cells during sorting (Figure S3A). Unsupervised clustering identified three clusters lacking TCR expression, which were excluded. The remainder was re-clustered into nine populations spanning naïve, effector, regulatory, and tissue-resident states (Figures 3A-C, S3B-D; Table S2). TCR transcripts were detected in over 90% of cells in every retained cluster, confirming these were αβT cells.

**Figure 3.**
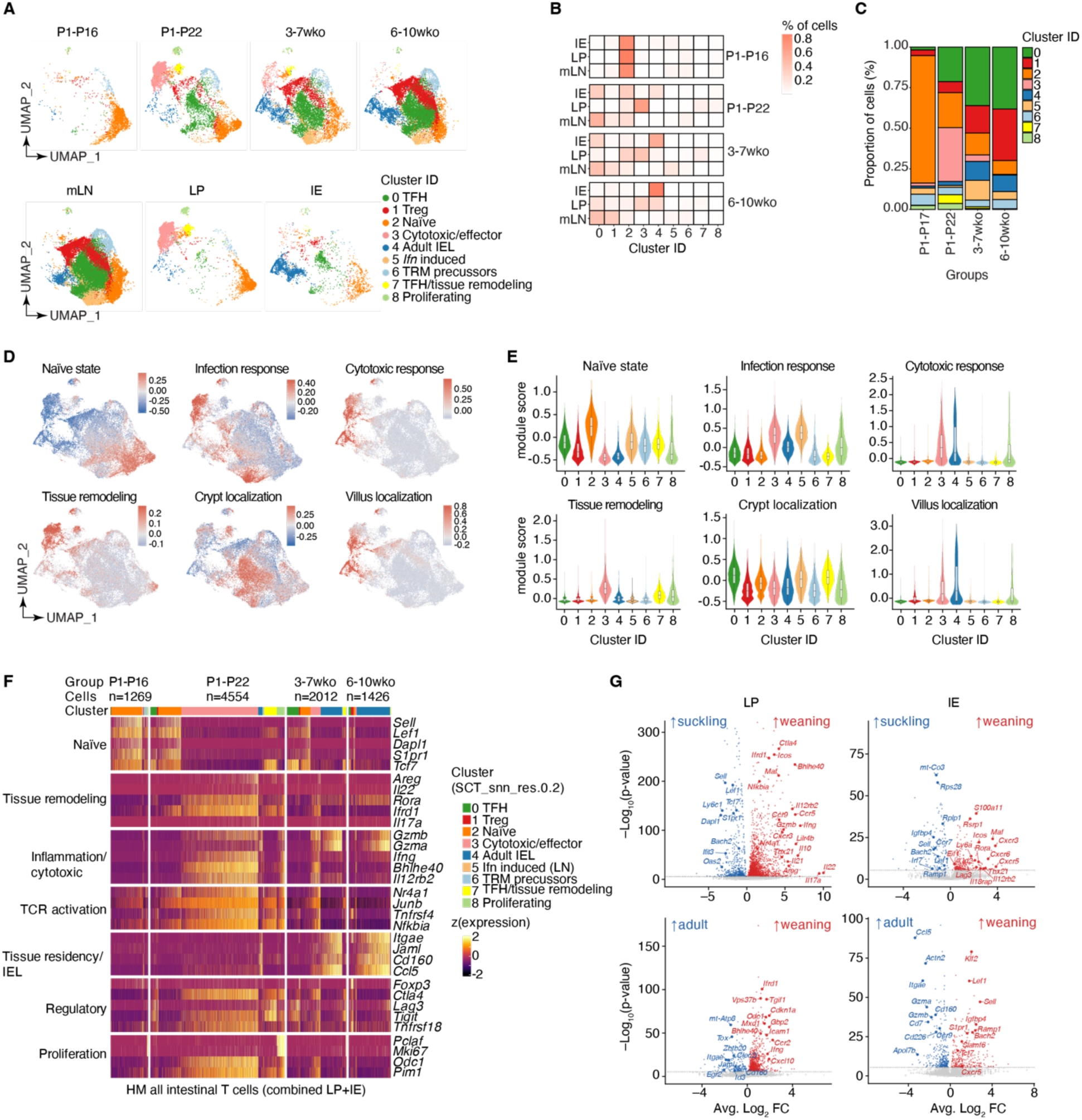
Weaning establishes a hybrid cytotoxic-regulatory program on lamina propria T lymphocytes. **(A)** UMAP of scRNA-seq data of timestamped CD4^+^ T cells across age and tissue. The top UMAPs were split by timestamping window, the bottom ones were separated by tissue: mLN, mesenteric lymph node; LP, lamina propria; IE, intestinal epithelium. **(B)** Distribution of T cells in clusters 0-8 by tissue compartment across timestamping windows. **(C)** Proportions of T cells in each cluster per timestamping window irrespective of tissue. **(D)** Gene module signatures visualized on scRNA-seq UMAPs of all CD4^+^ T cells. (E) Violin plots of module score signatures in (D) across clusters 0-8. **(F)** Heatmap of selected genes and transcriptional programs of LP and IE CD4^+^ T cells resolved per cluster. The number of cells in each time window is indicated above. **(G)** Volcano plots of differentially expressed genes comparing lamina propria (LP) and intestinal epithelium (IE) CD4^+^ T cells across ages. For adult samples, data from 3-7 wko and 6-10 wko were pooled, suckling is P1-P16, weaning is P1-P22.

The cluster distribution shifted sharply with age (Figure 3A-C). At P1-P16, cells were almost entirely naïve (cluster 2, marked by *Sell*, *Klf2*, and *Lef1*) regardless of tissue, even though our sorting enriched for CD62L^-/low^ T cells. At both weaning timepoints (P1-P22 and 3-7wko), naïve T cells were largely replaced by two LP-restricted populations: a cytotoxic/effector cluster (cluster 3, marked by the expression of *Ifng* and *Gzmb*) and a TFH/tissue-remodeling cluster (cluster 7, marked by *Bcl6*, *Cxcr5* and *Areg*). By 6-10wko, both compartments had converged on tissue-resident states (cluster 4, marked by *Itgae* and *Ccl5*) characteristic of adult intraepithelial lymphocytes (IELs), and the weaning-associated clusters had mostly resolved. T cells in cluster 3 displayed a hybrid signature, co-expressing cytotoxic/effector genes alongside immunoregulatory genes (*Ctla4* and *Tigit*) and, in a subset, a Tr1-like profile (*Tbx21*, *Il10*) (Figures 3F and S3B,C), suggesting that weaning can induce concurrent effector and regulatory programs within a single population instead of separate lineages or states.

Gene module scores (Table S3) further supported this progression from a naïve to resident state via an inflammatory-like route (Figure 3D-E). Infection- and cytotoxic-response scores rose specifically at weaning in LP-confined cluster 3. Overlaying crypt- and villus-localization signatures^26^ further distinguished the two weaning clusters: cluster 3 scored towards villus and cluster 7 towards crypt, while 6-10wko T cells scored almost uniformly for villus genes. Projection of our timestamped T cell clusters onto spatial transcriptomics of the adult small intestine (10x Visium HD ST dataset, 10X Genomics; see Methods) located suckling T cells preferentially by the crypts and weaning or adult T cells along the villus (Figure S3E), consistent with our imaging data (See Figure 2D).

The hybrid weaning state paired cytotoxic or effector genes with transcriptional regulators of NF-κB signaling and cell cycle exit, including *Mxd1* and *Ifrd1* (Figures 3F-G and S3C-D). This pairing may reflect a T cell-intrinsic mechanism to limit inflammation even as cytotoxic potential is acquired, consistent with the resolution of both programs by the adult steady state, when tissue-residency genes predominate (Figure 3F-G). Weaning and adult T cells also upregulated the gut-homing receptor *Ccr9* relative to their suckling counterparts, which parallels the increased expression of *Ccl25* in adult enterocytes (See Figure 1H-I). In summary, the naïve T cells seeding the gut at suckling acquire a hybrid cytotoxic-regulatory state specifically in the lamina propria at weaning that resolved into tissue-resident populations by adulthood. By weaning, LP T cells resembled the transcriptional state that IE T cells reached later in life (Figure 3G). The transcriptional changes accompanying weaning resembled those previously described in intestinal CD4^+^ T cells from mice infected with enteric pathogens^27^. We propose that weaning induces a transient state of physiological inflammation accompanied by mechanisms that may preserve tissue integrity.

### The cytotoxic-like program on lamina propria weaning T cells is regionally imprinted

Granzyme B (Gzmb) marked the cytotoxic arm of the hybrid transcriptional state that lamina propria CD4^+^ T cells acquired at weaning. To quantify this at the protein level, we used flow cytometry to measure Gzmb expression by timestamped CD4^+^ intestinal T cells across both pulse-chase schemes, together with cytokine or transcription factors that identify the major CD4^+^ T cell subsets typically found in the gut: IFNγ (TH1), Rorγt (TH17), and Foxp3 (Treg).

The proportion of timestamped Gzmb^+^ T cells peaked at weaning in the SILP of SPF mice in both birth-pulse (P1-22) and 4-week-windows (3-7wko) schemes (Figure 4A). In the epithelium, the rise in Gzmb^+^ and IFNγ^+^ T cells was instead delayed until adulthood, corroborating our gene expression analysis (see Figure 3G). Within the birth-timestamped cohort, Foxp3^+^ cells were most frequent shortly after birth (P1-P7) in both LP and IE (Figures 4A and S4A). We then compared Foxp3 and Gzmb expression between timestamped Tomato^+^ and non-timestamped Tomato^-^ CD4^+^ T cells at P1-P7 and P1-P56. At P1-P7, the frequency of Foxp3^+^ cells was higher among Tomato^+^ than Tomato^-^ T cells. Conversely, Gzmb-expressing T cells were enriched among timestamped T cells in P1-P56 mice (Figure S4A). Thus, the composition of the neonatal cohort shifted from regulatory to effector. The decrease in Treg cells among timestamped T cells could reflect limited survival, faster turnover or loss of *Foxp3* expression with differentiation into effector fates^28,29^, either of which would result in less Treg cells in adults. In both pulse schemes, timestamped IFNγ^+^ T cells were abundant in the LP at weaning and in adulthood but peaked in the IE only in adulthood. In contrast, Rorγt^+^ TH17 cells, though less abundant, peaked at weaning in both the LP and IE (Figures 4A and S4B-C). Together, these data reinforced highly compartmentalized T cell responses across tissue compartments and ages.

**Figure 4.**
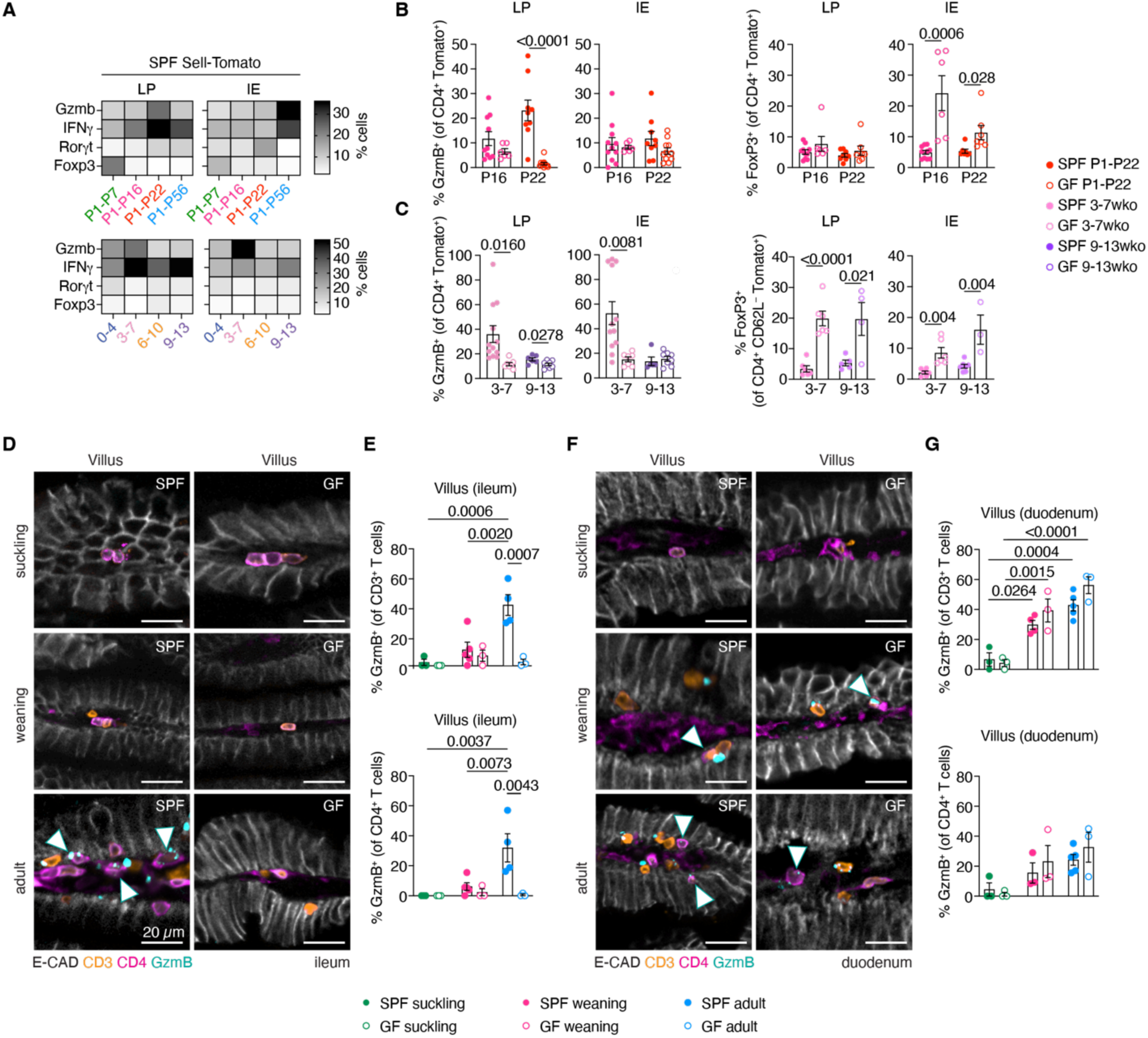
The cytotoxic-like program on lamina propria weaning T cells is regionally imprinted. **(A)** Heatmaps of flow cytometry analyses showing fraction of Gzmb^+^, IFNγ^+^, Rorγt^+^ and Foxp3^+^ CD4^+^ T cells across timestamping windows in SPF mice. (**B-C**) Flow cytometry analysis of the indicated populations of SPF and GF mice. (*n* = 5-12 mice/age). SPF mice used in the heatmap in (A) are shown here individually for comparison. **(D–G)** Representative sagittal sections of the ileal (D) and duodenal villus from SPF (left) and GF (right) mice at suckling (P7), weaning (P18), and adult (8 wko) stages. Quantification of Gzmb^+^ T cells (top) and Gzmb^+^ CD4^+^ T cells (bottom) in the ileum (E) and duodenum (G). (n = 3-5 mice/age). Bar plots show the mean ± SEM; each symbol represents one mouse (B-C, E and G). *P* values were calculated by unpaired Student’s t-tests comparing SPF and GF mice per timepoint in (B-C) and a one-way ANOVA with Tukey’s multiple comparisons (E, G).

The concomitant increase in Gzmb^+^ and IFNγ^+^ T cells at weaning prompted us to test whether this accumulation was dependent on the microbiota. Gzmb expression by timestamped CD4^+^ T cells was significantly higher in SPF mice than in GF mice at weaning (P1-22), especially in the SI LP, whereas in the IE it remained low in both (Figure 4B). This supports the notion that the SI LP acquires an effector phenotype before the SI IE. In adult mice, CD4^+^ T cells timestamped at weaning again showed high proportions of Gzmb-expressing CD4^+^ T cells in both the LP and IE, exceeding those in GF mice. By 13 weeks of age, these proportions were decreased and comparable between SPF and GF mice (Figure 4C). IFNγ^+^ T cells were consistently more frequent in SPF than in GF mice (Figure S4B-C), in agreement with previous reports^5,9^. GF mice displayed an increased frequency of Treg cells, which emerged in the epithelium during suckling and persisted in adulthood, but appeared in the LP only after weaning (Figure 4B-C). However, only the emergence of Gzmb^+^ CD4^+^ T cells was specific to weaning, consistent with a transient program of microbiota-induced physiological inflammation.

Exposure to microbes varies along the small intestinal epithelium, where crypts are protected by mucus and antimicrobial peptides from neighboring Paneth cells^30^. Paralleling these differences in microbial load, Gzmb-expressing T cells preferentially localized to the villi of SPF mice from weaning onward (Figure 4D-G). In the ileum, Gzmb^+^ T cells persisted into adulthood but were absent in GF mice, consistent with our flow cytometry analyses and confirming their dependence on the microbiota (Figures 4D-E). In contrast, Gzmb^+^ T cells appeared in duodenal villi of both SPF and GF mice at comparable levels around weaning (Figures 4F-G), suggesting that dietary stimulation may influence crypt-to-villus migration in the proximal intestine. Together, these data also point to another layer of compartmentalization, between the duodenum and ileum, that is initiated at weaning. Consistent with this, in published matched single-cell and spatial transcriptomic data^19^, *Gzmb* expression was enriched in villus T cells of adult SPF mice, whereas *Foxp3* expression peaked lower along the crypt-villus axis (Figure S4D). Our IBEX imaging of adult SPF ileum further supported this as Gzmb^+^ and Foxp3^+^ CD4^+^ T cells occupied the villi and crypts, respectively (Figure S4E). These data point to spatially segregated T cell responses across intestinal regions and along the crypt-villus axis in which T cells arriving at weaning adopt a transcriptional program modulated by the local tissue environment in an age- and region-specific manner.

### Weaning is marked by compartmentalized clonal expansions in the small intestinal lamina propria

Next, we investigated whether age-, layer-, and region-specific programs are accompanied by selective clonal expansion and how clonal dynamics unfold during postnatal development. Using key windows from both timestamping schemes (Figure 5A), we profiled the clonal architecture of intestinal CD4^+^ T cells across tissue compartments using plate-based and 10X Genomics scTCR-seq.

**Figure 5.**
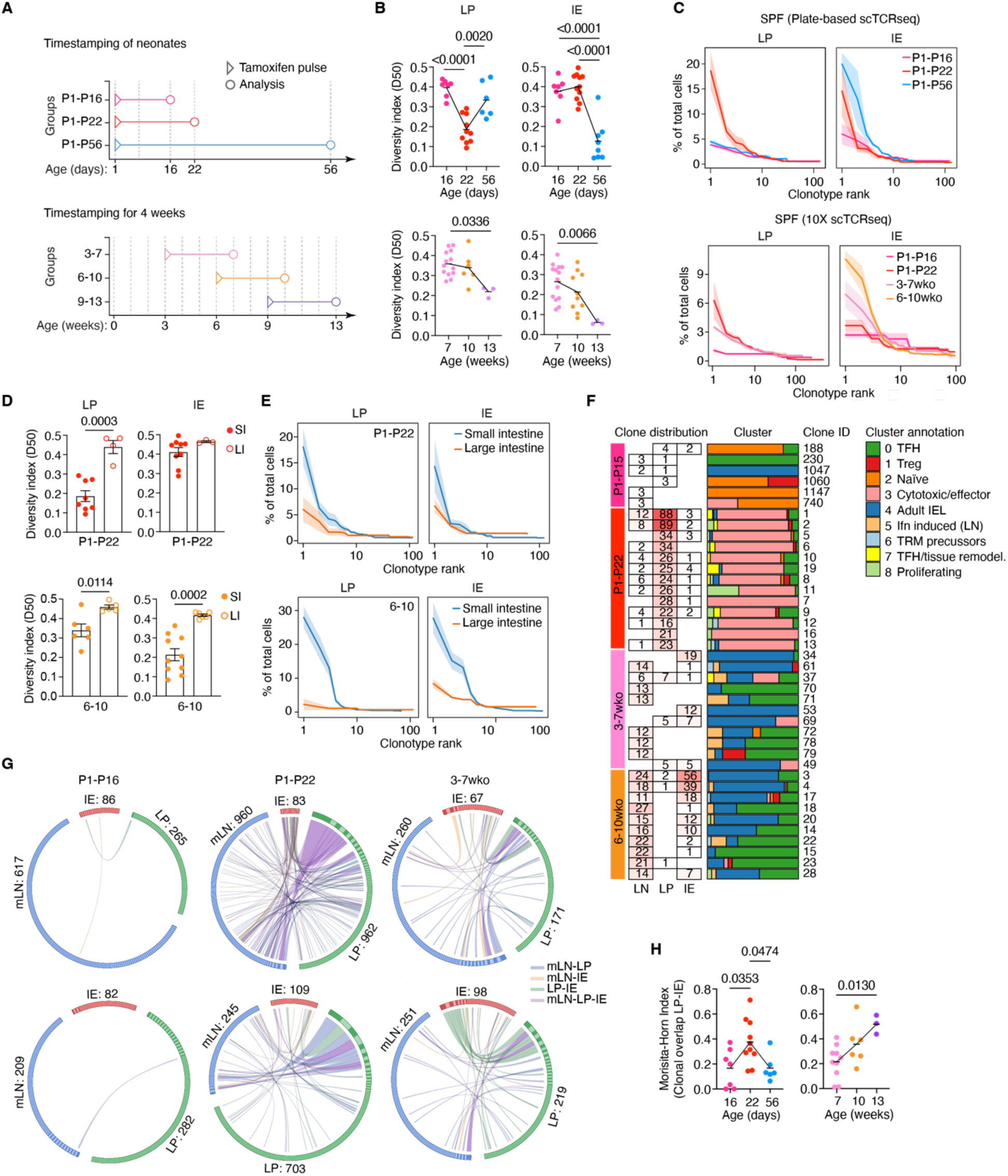
Weaning is marked by compartmentalized clonal expansions in the small intestinal lamina propria. **(A)** Timestamping windows for scTCR-sequencing in Sell-Tomato mice. **(B-E)** TCR clonal analyses of timestamped intestinal CD4^+^ T lymphocytes. TCR diversity was determined by the diversity index 50 (D50) in the indicated timestamping windows in (B) and (D). Clonotype-ranking analysis to show clonal dominance among total cells per mouse is in C and E. Curves and shaded areas show the mean and SEM, respectively, of all mice included. (n=3-14 mice/age). **(F)** Top expanded TCR clones in each timestamping window mapped across tissue compartment and transcriptional cluster of origin (10X scTCR-seq data). **(G)** Circos plots visualizing clonal overlap across tissue compartments (2 mice/age group). Arcs indicate tissue. Size of the slices within arcs indicates size of the clones. Links indicate tissue overlap between clones. **(H)** Clonal overlap between LP and IE determined by the Morisita-horn index (MHI) (n=3-10 mice/age group). *P* values were calculated by one-way ANOVA with Tukey’s multiple comparison test for groups larger than two or an unpaired Student’s t-test for groups in D.

T cells timestamped at birth and analyzed at P16 were clonally diverse in both the SI LP and IE as measured by the diversity index 50 (D50; Figure 5B). By weaning (P1-P22), the birth cohort had undergone clonal expansion specifically in the lamina propria, whereas in the IE it remained clonally diverse (Figure 5B). In adulthood (P1-P56), the pattern reversed as T cells were more expanded in the IE than LP. The 4-week-window scheme supported this asymmetry as T cells timestamped at weaning or in adulthood remained relatively diverse in the LP but were clonally expanded in the IE, with expansion increasing in both compartments with age (Figure 5B). These data indicate that clonal diversity narrows with age and selected clones can establish long-term residence in the epithelium, while lamina propria clones appear to be replaced at a higher rate in adulthood.

Clonotype ranking, from most to least abundant clone within each group, showed that 6-12% of intestinal T cells belonged to the same clone in the weaning lamina propria, where expansion first occurred (Figure 5C). Dominance was highest among adult IELs, where a given clone accounted for ∼20% of the population in some mice, consistent with their low TCR diversity 50 index. T cells isolated from the large intestine and mLN remained diverse throughout life (Figures 5D-E and S5A-B), indicating that clonal expansion in SPF mice is restricted to the small intestine.

To relate clonal expansion to fate, we mapped the most expanded clones identified in each age group from the 10X scTCR-seq dataset to their transcriptional clusters. Clones at weaning belonged predominantly to the cytotoxic/effector cluster, with some overlap with TFH and Treg clusters (Figure 5F). In contrast, adult clones mapped to both the IEL and TFH clusters. Clones from the Treg cluster in our dataset were not expanded (Figure S5C; Table S5). A subset of clones that were timestamped at weaning but expanded in adulthood spanned the IEL and cytotoxic/effector weaning clusters (Figure 5F), suggesting that clones that expanded during weaning can persist and become resident in the epithelium. Expanded weaning clones were mostly confined to the LP, whereas adult clones were distributed across all three compartments. Circos plots linked compartments by shared clonal identity (two mice per age group). At P16, before expansion occurs, almost no clones overlapped between compartments. By weaning and adulthood, clones were shared extensively between lamina propria, epithelium, and, to a lesser extent, mLN (Figure 5G). Analysis using Morisita-Horn index confirmed that, among T cells timestamped at birth, LP and IE overlap was already high at weaning, whereas cells timestamped at weaning or in adulthood showed the highest LP-IE overlap in the oldest (9-13wko) cohort (Figure 5H). More clones were shared between mLN and LP than between mLN and IE at both weaning and adulthood (Figure S5C), consistent with the lamina propria, but not the epithelium, continuing to exchange clones with the recirculating pool.

Finally, we next asked to what extent the microbiota could modulate the TCR repertoire of intestinal CD4^+^ T cells during postnatal development. In GF mice, T cells timestamped at birth did not undergo clonal expansion in the LP at weaning. However, T cells that entered the gut after weaning, expanded equally in GF and SPF mice (Figure S5D). Clonal overlap between LP and IE was low in GF mice at weaning but rose in adulthood, matching if not exceeding the overlap in SPF mice (Figure S5E). Thus, at weaning, clonal expansion of T cells timestamped at birth requires the microbiota, whereas expansion of T cells timestamped in adulthood occurs in both SPF and GF mice. These data suggest a model for how the intestinal T cell repertoire is selected and stabilized as it encounters the microbiota during the suckling-to-weaning transition.

### Expanded weaning clones react to early-life microbiota

The increased proportion of granzyme B and IFNγ-expressing cells, together with clonal expansions at weaning in SPF, but not GF mice (See Figures 4 and 5), suggested that weaning-expanded clones recognize the microbiota, which amplifies and diversifies at this time. To profile their antigen reactivity, we reconstructed the top expanded TCRs in NFAT-GFP T cell hybridomas^31^ and screened them *in vitro* against intestinal luminal content from suckling, weaning, and adult SPF mice, each enriched in age-specific microbiota (Figure 6A-B). All nine weaning clones reacted to suckling luminal content, five also reacted to weaning content, and one reacted to content from all three stages. In contrast, only one adult clone reacted to the weaning content (Figure 6B). With one exception, no weaning clone reacted to GF luminal content or to the food antigens tested (Figure S6A), suggesting that TCR reactivity among expanded weaning clones is focused on the commensal microbiota rather than diet. Among the adult clones tested, two reacted to food-derived antigens (Figure S6A). Because we screened only the top expanded clones from each age group against crude antigenic mixtures, some reactivities were likely missed. The low reactivity to adult SPF content may also reflect the dilution of putative ligands in a more antigenically diverse mixture rather than an age-specific difference. Nonetheless, these data suggest that weaning clones preferentially react to early-life microbiota.

**Figure 6.**
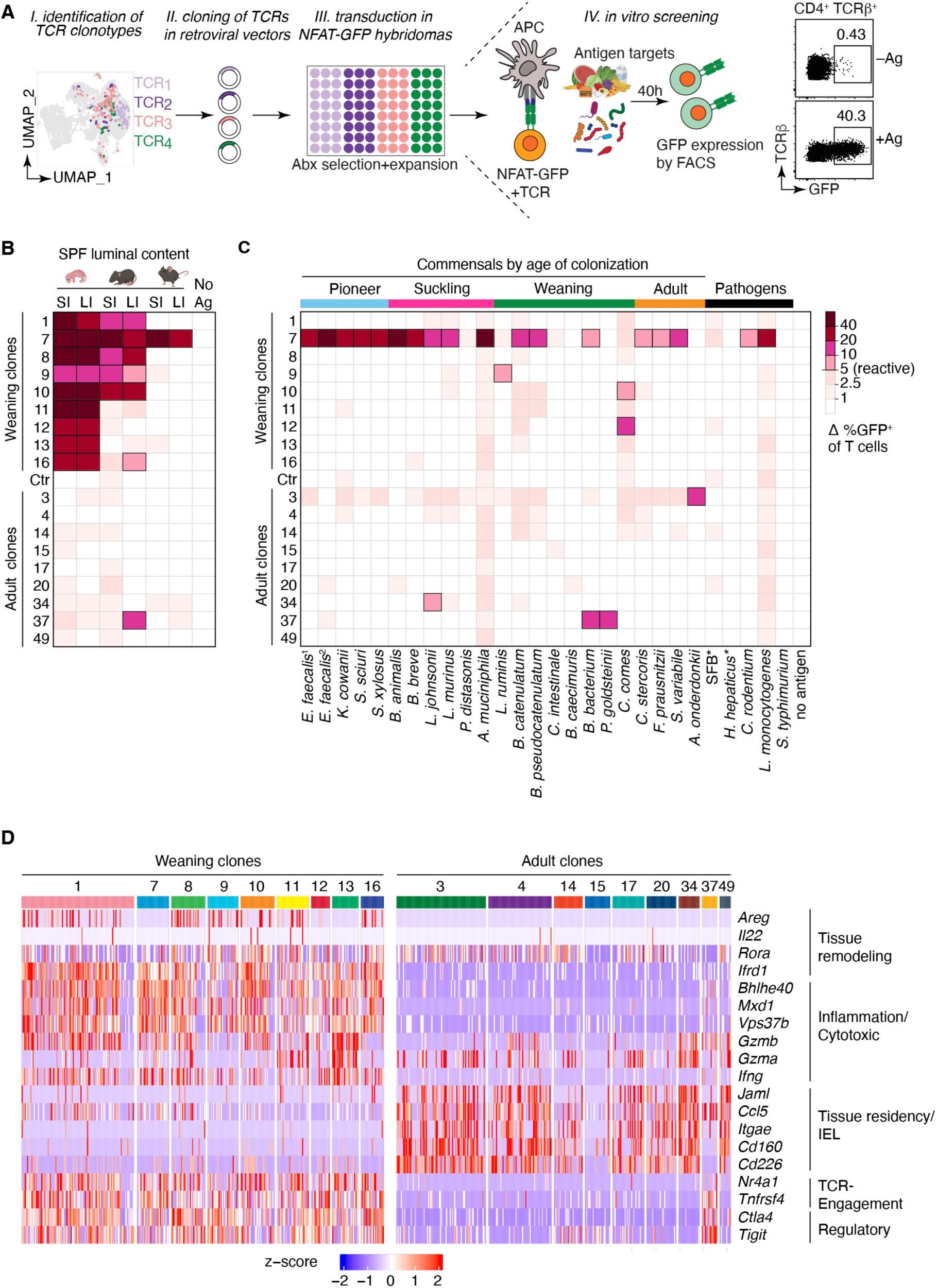
Expanded weaning clones react to early-life microbiota. **(A)** Schematic of the hybridoma-based antigen-reactivity screen. **(B-C)** Heatmaps show reactivity of the indicated expanded-weaning- or adult clones to the indicated antigens measured by frequency of GFP^+^ T cells. Luminal content from the small (SI) or large (LI) intestine of suckling, weaning, and adult SPF mice were tested (B). Reactivity to the indicated individual commensal bacterial strains, ordered by their approximate age of first colonization (pioneer, suckling, weaning, adult) or by pathogen status (C). No antigen, no antigen stimulation; Ctr, control TCR with mispaired TCRαβ; ^1^PedsCom strain; ^2^Oligo-MM^12^ strain; *Pathobionts. **(D)** Heatmap of gene module expression (z-score) of the indicated genes in expanded-weaning and adult clones grouped into the indicated gene signatures.

To identify reactivity to bacterial species, we screened defined mouse and human commensal consortia whose genera and species are also represented in the microbiota of our facility^9–11,32^. After classifying species by their age of colonization (see Methods, Table S6), four of nine weaning clones and three of nine adult clones reacted to at least one species (Figures 6C and S6C). Most weaning clones reacted to a single species, whereas one “generalist” (clone 7) reacted to species from every colonization stage and to enteric pathogens, to varying degrees.

Two adult clones timestamped at weaning (clones 34 and 37) reacted to suckling and weaning colonizers whereas adult clone 3 timestamped in adulthood (See Figure 5F) reacted to an adult colonizer and to additional species from the human consortium screen (Figures 6C and S6C). Because food preparations may contain residual bacterial protein, the food reactivity of the commensal-reactive clones 7 and 37 may reflect bacterial rather than dietary antigens. The chow-only reactivity of adult clone 20 more likely reflects a dietary antigen, although a bacterial origin cannot be excluded. Because the panels do not cover all the microbial species and food proteins to which SPF mice are exposed, additional reactivities cannot be excluded. In the clones tested, commensal reactivity was MHC-II-dependent, as anti-MHC-II blocking reduced it to baseline (Figure S6D-E).

Reactive weaning clones were expanded mainly in the cytotoxic/effector cluster, with some cells in the TFH and Treg clusters (See Figure 5F). Gene expression differed between weaning and adult clones (Figure 6D). Most adult clones, which expanded in the IE, expressed tissue-residency/IEL genes (*Itgae*, *Cd160*). Weaning clones, which expanded in the LP, instead expressed genes associated with TCR engagement (*Nr4a1*), inflammation/cytotoxicity (*Bhlhe40*, *Ifng*), and regulation (*Ctla4*, *Tigit*), without resolving into one dominant signature. For example, most cells of generalist clone 7 had a strong inflammatory/cytotoxic signature (*Gzmb*, *Ifng*) with no expression of *Areg* or *Rora*, whereas *Lactobacillus*-reactive clone 9 showed the inverse, and cells of clone 1 displayed all three signatures (tissue remodeling, inflammatory/cytotoxic, and regulatory). Because specificity, age, and tissue of expansion are intertwined in this dataset and only a handful of clones were examined, these observations do not imply that antigen specificity dictates a clone’s functional state. They illustrate the spectrum of states possible. Together, these data show that microbiota-dependent clonal expansion at weaning includes commensal-reactive clones that coexist with a hybrid cytotoxic-regulatory state.

## Discussion

Following T cells across postnatal development revealed that weaning transiently imprints a hybrid cytotoxic-regulatory state on lamina propria CD4⁺ T cells that coincided with microbiota-dependent clonal expansions, many of which recognized early-life commensals. Weaning is accompanied by inflammatory signals but proceeds without overt tissue damage, suggesting that the regulatory features of this state may help temper inflammation.

Our data suggest a two-gate model of intestinal T cell entry, in which CD62L-assisted entry during suckling gives way to integrin-dependent entry after weaning. During suckling, T cells express CD62L, coinciding with increased MAdCAM-1^+^ vessel density. Beyond its canonical receptor, integrin α4β7, whose expression by T cells is stable across ages, MAdCAM-1 can also bind CD62L through its mucin-like domain^33^. This suggests that CD62L could assist in early tethering and naïve T cell entry into the intestinal mucosa. MAdCAM-1^+^ vessels direct T cells toward the crypt in the adult intestine after allogeneic transplantation^34^, in agreement with T cells entering near the crypt by default during development and redistributing to the villus with age. After weaning, when CD62L is lost, entry may rely on the classical α4β7-MAdCAM-1 axis, consistent with α4β7 being required for T cell accumulation in both the neonatal^4^ and adult gut^35^.

The location of T cells depends on a microenvironment that changes with age. Epithelial expression of *Il15*, *Ccl25*, and MHC-II-related genes increased with age, so a neonatal environment low in these cues may retain incoming T cells near the crypt. Blimp-1 (*Prdm1)*, a transcriptional repressor that is highly expressed in the neonatal epithelium^22^ and can suppress MHC-II in immune cells^20,21^, was inversely correlated with MHC-II expression in epithelial cells across ages. This suggests that Blimp-1 may limit epithelial MHC-II and engagement with incoming CD4^+^ T cells during suckling. Around weaning, when intestinal epithelial cells upregulate MHC-II expression in a microbiota-dependent manner, the epithelium may be licensed to present antigens to CD4^+^ T cells, allowing T cells to differentiate into effector subsets typical of adult mice^36–41^.

Our timestamping data show that weaning drives a substantial influx of T cells to the gut but also that T cells seed the intestine earlier during suckling. Despite their early arrival and long-term persistence in the tissue, timestamped T cells underwent their most pronounced transcriptional changes in the lamina propria at weaning, adopting a hybrid state that combined both cytotoxic and regulatory features. While a cytotoxic-like program parallels an increase in pro-inflammatory signals (TNF-α and IFN-γ) in the tissue during weaning^5^, the simultaneous emergence of regulatory profiles may restrain these signals and help preserve tissue integrity. We propose that this balance may hinge on the microbial load, as IFN-γ⁺ T cells were more frequent in SPF intestine while Foxp3⁺ Treg cells were more frequent in GF mice. Gzmb-expressing CD4^+^ T cells were specifically enriched at weaning and accumulated preferentially in the villus, where effector CD8^+^Gzmb^+^ LCMV-specific IELs also reside^26^. A Gzmb-associated phenotype has also been shown to be triggered by diet^42^, and a diet-driven, microbiota-independent Gzmb/Tbet^+^ Tr1 population emerges in the small intestine around weaning^43^. Consistent with distinct roles for diet and microbiota, the weaning-associated Gzmb program was microbiota-independent in the duodenum, where dietary antigens are concentrated, but microbiota-dependent in the ileum, where microbial load is higher. Gzmb expression may thus emerge in response to signals from both diet and microbiota, allowing regionalized responses in the intestinal mucosa.

At weaning, clonal expansion of intestinal CD4^+^ T cells was restricted to the small intestinal lamina propria and absent in GF mice. Screening these expanded clones for antigen reactivity suggested early-life microbiota as a driver of expansion. Clones ranged from generalists, reacting to multiple bacterial species, to specialists, reacting to a single species. We detected little reactivity among the adult clones tested. Although we cannot exclude age-related differences in reactivity, this may reflect the lower relative abundance of their ligands in the more diverse adult luminal content. Alternatively, these mostly epithelium-derived clones may require a higher threshold for TCR activation, as IELs typically do^44,45^, which could mask reactivity under the conditions of our assay. Larger clone panels will help assess the relative contribution of antigen specificity and tissue context in determining the cell state.

Our weaning-expanded clones predominantly displayed an effector phenotype, with some cells of the same clonotype also present in the TFH and Treg clusters. This is consistent with the presence of both effector and Treg microbiota-reactive clones in adult GF mice two weeks after colonization with a defined human microbial community^10^. This model of colonization exposes the intestinal immune system to an abrupt burst of new antigens, like the weaning period, allowing the establishment of an effector compartment at the intestinal mucosa. In contrast, microbiota-reactive clones isolated from mice vertically colonized from birth with defined mouse or human consortia were predominantly Treg cells, suggesting that commensal reactivity is biased toward a Treg fate^46,47^. Defined consortia may lack the diversity and antigenicity of the SPF microbiota, favoring Treg accumulation, as in GF mice, which do not experience a weaning reaction.

Our fate-mapping did not capture expanded Treg cells and cannot distinguish whether timestamped Gzmb^+^ T cells differentiate directly from naïve precursors or pass through a transient Foxp3^+^ regulatory state. Foxp3 instability has been described^28^, including the differentiation of Treg cells into CD4^+^CD8αα^+^ T cells specifically at the intestinal epithelium, in a microbiota-dependent manner^29^, which makes a transient Treg intermediate a plausible route to the effector program we observed in timestamped T cells. Alternatively, timestamped Treg cells may have been outcompeted by effector cells because differentiation at weaning is biased toward effector fates.

Beyond cell state, weaning-expanded clones were also shared between intestinal compartments. This clonal overlap, which began at weaning, suggests a route from the lamina propria to the epithelium, where most of the adult clones we screened resided. Overlap increased with age as epithelial clones expanded and lamina propria clones turned over, consistent with weaning expansions feeding the epithelial pool, although direct entry into the epithelium cannot be excluded. We have previously shown that Treg cells and conventional CD4⁺ T cells undergo a TCR-dependent bottleneck in which selected clones can expand and become epithelium residents, a process that depends on MHCI-II expression on epithelial cells^38^. The antigens driving these expansions may derive from the microbiota, diet, or both. In SPF mice, microbiota-dependent expansions at weaning point to a primary role for the microbiota, without precluding contributions from food or self-antigens. Consistent with a dietary contribution, food induces Tregs in the small intestine^42,48,49^. Likewise, T cells specific for the dietary antigen alpha-zein are enriched among Treg cells that also accumulate at weaning^50^, indicating that weaning supports diet-specific Treg induction in parallel. In GF mice, adult-timestamped T cells clonally expanded as in SPF mice, suggesting that dietary or self-antigens can drive clonal expansion without a microbiota, in agreement with food- and self-reactive TCRs driving peripheral Treg cell differentiation^51^. Because antigen class and TCR origin both influence Treg cell differentiation^51^, the antigens that select epithelial residents may also influence whether these clones adopt a regulatory or effector fate.

Together, these findings show that the intestinal T cell compartments are established in stages, with weaning as a specific window in which lamina propria CD4⁺ T cells acquire a hybrid cytotoxic-regulatory state, and commensal-reactive clones expand. We propose that the concomitant cytotoxic-regulatory features of this state may help the gut engage the rapid microbial expansion that occurs at weaning while limiting damage. By seeding a spatially organized pool of persistent T cells, these early-life encounters may help establish immune homeostasis in the adult gut, alongside clonal expansions during adulthood that do not depend on the microbiota.

## RESOURCE AVAILABILITY

### Lead Contact

Further information and requests for resources and reagents should be directed to and will be fulfilled by the Lead Contacts, Daniel Mucida or Angelina M. Bilate

### Materials Availability

All animal strains used in this study are available from The Jackson Laboratory or were provided by the indicated investigators. This study did not generate any new unique reagents.

## ACKNOWLEDGEMENTS

We are grateful to A. Rogoz and L. Bianco for exceptional animal care, mouse colony management, and genotyping; RU Genomics core for assistance with sequencing; K. Gordon and J. P. Truman for sorting; Y. Khan, G. Scrivanti, and additional Rockefeller University employees for continuous assistance. We thank D. Littman for providing *H. hepaticus*, M. Fischbach for providing the hCom2 consortium, M. Silverman for providing the PedsCom consortium, and K. McCoy for providing the OligoMM^12^ consortium. We thank the NIH Tetramer Core Facility for providing the CD1d tetramer. We thank all members of the Mucida, Victora (Rockefeller), Fuchs (Rockefeller), and Lafaille (NYU) laboratories for fruitful discussions. This work was supported by NIH R01DK093674, R01DK113375, Chan Zuckerberg Initiative (CZI) science and Food Allergy FARE/FASI Consortium and The Howard Hughes Medical Institute (D.M., E.F.). M.S was a Rockefeller University Women & Science and Boehringer Ingelheim Fonds graduate fellow. G.L.d.R. was initially supported by a scholarship from Fundação Maria Emília (Brazil). This work also benefited from discussions during the 2024 program “Interactions and Co-evolution between Viruses and Immune Systems” at the Kavli Institute for Theoretical physics (KITP), which is supported by National Science Foundation grants PHY-2309135, and the Gordon and Betty Moore Foundation Grant No. 2919.02.

## AUTHOR CONTRIBUTIONS

A.M.B. and D.M. conceived the study. A.M.B. and M.S initiated the study and designed experiments. A.M.B., M.S and D.M. wrote the manuscript. M.S., R.R.G., G.L.d.R., R.P., and A.M.B. performed and analyzed experiments. I.M. performed initial bioinformatics analysis, and T.B.R.C. performed and supervised bioinformatics analyses and assisted with interpretation of sequencing data. A.T. M.Sa. M.M.C. performed NFAT-GFP hybridoma experiments. Y.S.Y. and R.P. assisted with plate-based scTCR-seq analysis. B.S.R. performed 2-photon microscopy. A.M.B. and D.M. supervised the research. E.F., A.M.B., and D.M. acquired funding. All authors edited the manuscript.

## DECLARATION OF INTERESTS

The authors declare no competing interests

### Declaration of generative AI and AI-assisted

During the preparation of this work, the authors used Claude, Claude Code, and Claude Science (Anthropic) to assist with data analysis, literature search, figure plotting, and text editing. After using these tools, the authors reviewed and edited the content as needed and take full responsibility for the content of the published article.

### Notice of pre-existing conditions, requirements, and licenses for article submission

The article submitted together with this notice is subject to the Immediate Access to Research policy of the Howard Hughes Medical Institute (“HHMI”). In accordance with this policy: (i) a preprint of this article either has been, or will be, deposited on a preprint server under a Creative Commons Attribution 4.0 International (CC BY 4.0) license and (ii) an additional author-published revised version of this article incorporating peer review feedback and/or new results or analysis either has been, or prior to journal publication will be, deposited on a preprint server under a CC BY 4.0 license. In addition, a non-exclusive CC BY 4.0 license to this article has been granted to the public and HHMI has a sublicensable, non-exclusive license to this article. THIS ARTICLE IS SUBMITTED FOR REVIEW AND ACCEPTANCE SUBJECT TO THESE PRE-EXISTING CONDITIONS, REQUIREMENTS AND LICENSES. If you have any concerns with any of these pre-existing conditions, requirements or licenses, please contact the corresponding author immediately.

## REFERENCES

1. Corral, D., Ansaldo, E., Delaleu, J., Pichler, A.C., Kabat, J., Oguz, C., Teijeiro, A., Yong, D., Abid, M., Rivera, C.A., et al. (2025). Mammary intraepithelial lymphocytes promote lactogenesis and offspring fitness. Cell 188, 1662–1680.e24. 10.1016/j.cell.2025.01.028.

2. Waizman, D.A., and Wang, A. (2025). A physiological perspective on neuroimmune tissue innervation. Trends Immunol. 46, 508–511. 10.1016/j.it.2025.05.003.

3. Florsheim, E.B., Sullivan, Z.A., Khoury-Hanold, W., and Medzhitov, R. (2021). Food allergy as a biological food quality control system. Cell 184, 1440–1454. 10.1016/j.cell.2020.12.007.

4. Torow, N., Yu, K., Hassani, K., Freitag, J., Schulz, O., Basic, M., Brennecke, A., Sparwasser, T., Wagner, N., Bleich, A., et al. (2015). Active suppression of intestinal CD4+TCRαβ+ T-lymphocyte maturation during the postnatal period. Nat. Commun. 6, 7725. 10.1038/ncomms8725.

5. Nabhani, Z.A., Dulauroy, S., Marques, R., Cousu, C., Bounny, S.A., Déjardin, F., Sparwasser, T., Bérard, M., Cerf-Bensussan, N., and Eberl, G. (2019). A Weaning Reaction to Microbiota Is Required for Resistance to Immunopathologies in the Adult. Immunity 50, 1276–1288.e5. 10.1016/j.immuni.2019.02.014.

6. Lathrop, S.K., Bloom, S.M., Rao, S.M., Nutsch, K., Lio, C.-W., Santacruz, N., Peterson, D.A., Stappenbeck, T.S., and Hsieh, C.-S. (2011). Peripheral education of the immune system by colonic commensal microbiota. Nature 478, 250–254. 10.1038/nature10434.

7. Yang, Y., Torchinsky, M.B., Gobert, M., Xiong, H., Xu, M., Linehan, J.L., Alonzo, F., Ng, C., Chen, A., Lin, X., et al. (2014). Focused Specificity of Intestinal Th17 Cells towards Commensal Bacterial Antigens. Nature 510, 152–156. 10.1038/nature13279.

8. Ansaldo, E., Slayden, L.C., Ching, K.L., Koch, M.A., Wolf, N.K., Plichta, D.R., Brown, E.M., Graham, D.B., Xavier, R.J., Moon, J.J., et al. (2019). Akkermansia muciniphila induces intestinal adaptive immune responses during homeostasis. Science 364, 1179–1184. 10.1126/science.aaw7479.

9. Bousbaine, D., Fisch, L.I., London, M., Bhagchandani, P., Castro, T.B.R. de, Mimee, M., Olesen, S., Reis, B.S., VanInsberghe, D., Bortolatto, J., et al. (2022). A conserved Bacteroidetes antigen induces anti-inflammatory intestinal T lymphocytes. Science 377, 660–666. 10.1126/science.abg5645.

10. Nagashima, K., Zhao, A., Atabakhsh, K., Bae, M., Blum, J.E., Weakley, A., Jain, S., Meng, X., Cheng, A.G., Wang, M., et al. (2023). Mapping the T cell repertoire to a complex gut bacterial community. Nature 621, 162–170. 10.1038/s41586-023-06431-8.

11. Brugiroux, S., Beutler, M., Pfann, C., Garzetti, D., Ruscheweyh, H.-J., Ring, D., Diehl, M., Herp, S., Lötscher, Y., Hussain, S., et al. (2016). Genome-guided design of a defined mouse microbiota that confers colonization resistance against Salmonella enterica serovar Typhimurium. Nat. Microbiol. 2, 16215. 10.1038/nmicrobiol.2016.215.

12. Cheng, A.G., Ho, P.-Y., Aranda-Díaz, A., Jain, S., Yu, F.B., Meng, X., Wang, M., Iakiviak, M., Nagashima, K., Zhao, A., et al. (2022). Design, construction, and in vivo augmentation of a complex gut microbiome. Cell 185, 3617–3636.e19. 10.1016/j.cell.2022.08.003.

13. Afrizal, A., Jennings, S.A.V., Hitch, T.C.A., Riedel, T., Basic, M., Panyot, A., Treichel, N., Hager, F.T., Wong, E.O.-Y., Wolter, B., et al. (2022). Enhanced cultured diversity of the mouse gut microbiota enables custom-made synthetic communities. Cell Host Microbe 30, 1630–1645.e25. 10.1016/j.chom.2022.09.011.

14. Beura, L.K., Hamilton, S.E., Bi, K., Schenkel, J.M., Odumade, O.A., Casey, K.A., Thompson, E.A., Fraser, K.A., Rosato, P.C., Filali-Mouhim, A., et al. (2016). Normalizing the environment recapitulates adult human immune traits in laboratory mice. Nature 532, 512–516. 10.1038/nature17655.

15. Abolins, S., King, E.C., Lazarou, L., Weldon, L., Hughes, L., Drescher, P., Raynes, J.G., Hafalla, J.C.R., Viney, M.E., and Riley, E.M. (2017). The comparative immunology of wild and laboratory mice, Mus musculus domesticus. Nat. Commun. 8, 14811. 10.1038/ncomms14811.

16. Constantinides, M.G., and Belkaid, Y. (2021). Early-life imprinting of unconventional T cells and tissue homeostasis. Science 374, eabf0095–eabf0095. 10.1126/science.abf0095.

17. Ganal-Vonarburg, S.C., and Duerr, C.U. (2020). The interaction of intestinal microbiota and innate lymphoid cells in health and disease throughout life. Immunology 159, 39–51. 10.1111/imm.13138.

18. Radtke, A.J., Kandov, E., Lowekamp, B., Speranza, E., Chu, C.J., Gola, A., Thakur, N., Shih, R., Yao, L., Yaniv, Z.R., et al. (2020). IBEX: A versatile multiplex optical imaging approach for deep phenotyping and spatial analysis of cells in complex tissues. Proc. Natl. Acad. Sci. 117, 33455–33465. 10.1073/pnas.2018488117.

19. Niec, R.E., Chu, T., Schernthanner, M., Gur-Cohen, S., Hidalgo, L., Pasolli, H.A., Luckett, K.A., Wang, Z., Bhalla, S.R., Cambuli, F., et al. (2022). Lymphatics act as a signaling hub to regulate intestinal stem cell activity. Cell Stem Cell 29, 1067–1082.e18. 10.1016/j.stem.2022.05.007.

20. Yang, H., Qiu, Q., Gao, B., Kong, S., Lin, Z., and Fang, D. (2014). Hrd1-mediated BLIMP-1 ubiquitination promotes dendritic cell MHCII expression for CD4 T cell priming during inflammation. J. Exp. Med. 211, 2467–2479. 10.1084/jem.20140283.

21. Piskurich, J.F., Lin, K.-I., Lin, Y., Wang, Y., Ting, J.P.-Y., and Calame, K. (2000). BLIMP-1 mediates extinction of major histocompatibility class II transactivator expression in plasma cells. Nat. Immunol. 1, 526–532. 10.1038/82788.

22. Muncan, V., Heijmans, J., Krasinski, S.D., Büller, N.V., Wildenberg, M.E., Meisner, S., Radonjic, M., Stapleton, K.A., Lamers, W.H., Biemond, I., et al. (2011). Blimp1 regulates the transition of neonatal to adult intestinal epithelium. Nat. Commun. 2, 452. 10.1038/ncomms1463.

23. Merkenschlager, J., Finkin, S., Ramos, V., Kraft, J., Cipolla, M., Nowosad, C.R., Hartweger, H., Zhang, W., Olinares, P.D.B., Gazumyan, A., et al. (2021). Dynamic regulation of TFH selection during the germinal centre reaction. Nature 591, 458–463. 10.1038/s41586-021-03187-x.

24. Parsa, R., London, M., Castro, T.B.R. de, Reis, B., Amorie, J.B. des, Smith, J.G., and Mucida, D. (2022). Newly recruited intraepithelial Ly6A+CCR9+CD4+ T cells protect against enteric viral infection. Immunity 55, 1234–1249.e6. 10.1016/j.immuni.2022.05.001.

25. Min, B., McHugh, R., Sempowski, G.D., Mackall, C., Foucras, G., and Paul, W.E. (2003). Neonates Support Lymphopenia-Induced Proliferation. Immunity 18, 131–140. 10.1016/s1074-7613(02)00508-3.

26. Reina-Campos, M., Monell, A., Ferry, A., Luna, V., Cheung, K.P., Galletti, G., Scharping, N.E., Takehara, K.K., Quon, S., Challita, P.P., et al. (2025). Tissue-resident memory CD8 T cell diversity is spatiotemporally imprinted. Nature 639, 483–492. 10.1038/s41586-024-08466-x.

27. Kiner, E., Willie, E., Vijaykumar, B., Chowdhary, K., Schmutz, H., Chandler, J., Schnell, A., Thakore, P.I., LeGros, G., Mostafavi, S., et al. (2021). Gut CD4+ T cell phenotypes are a continuum molded by microbes, not by TH archetypes. Nat. Immunol. 22, 216–228. 10.1038/s41590-020-00836-7.

28. Zhou, X., Bailey-Bucktrout, S.L., Jeker, L.T., Penaranda, C., Martínez-Llordella, M., Ashby, M., Nakayama, M., Rosenthal, W., and Bluestone, J.A. (2009). Instability of the transcription factor Foxp3 leads to the generation of pathogenic memory T cells in vivo. Nat. Immunol. 10, 1000–1007. 10.1038/ni.1774.

29. Sujino, T., London, M., Konijnenburg, D.P.H. van, Rendon, T., Buch, T., Silva, H.M., Lafaille, J.J., Reis, B.S., and Mucida, D. (2016). Tissue adaptation of regulatory and intraepithelial CD4+ T cells controls gut inflammation. Science 352, 1581–1586. 10.1126/science.aaf3892.

30. Gehart, H., and Clevers, H. (2019). Tales from the crypt: new insights into intestinal stem cells. Nat. Rev. Gastroenterol. Hepatol. 16, 19–34. 10.1038/s41575-018-0081-y.

31. Parsa, R., Assis, H.C., Castro, T.B.R. de, Reis, G.L. dos, Sushil, A., Hartweger, H., Bilate, A.M., and Mucida, D. (2026). Tissue-specific clonal selection and differentiation of CD4^+^ T cells during infection. Nat. Immunol., 1–13. 10.1038/s41590-026-02451-4.

32. Lubin, J.-B., Green, J., Maddux, S., Denu, L., Duranova, T., Lanza, M., Wynosky-Dolfi, M., Flores, J.N., Grimes, L.P., Brodsky, I.E., et al. (2023). Arresting microbiome development limits immune system maturation and resistance to infection in mice. Cell Host Microbe 31, 554–570.e7. 10.1016/j.chom.2023.03.006.

33. Berg, E.L., McEvoy, L.M., Berlin, C., Bargatze, R.F., and Butcher, E.C. (1993). L-selectin-mediated lymphocyte rolling on MAdCAM-1. Nature 366, 695–698. 10.1038/366695a0.

34. Fu, Y.-Y., Egorova, A., Sobieski, C., Kuttiyara, J., Calafiore, M., Takashima, S., Clevers, H., and Hanash, A.M. (2019). T Cell Recruitment to the Intestinal Stem Cell Compartment Drives Immune-Mediated Intestinal Damage after Allogeneic Transplantation. Immunity 51, 90–103.e3. 10.1016/j.immuni.2019.06.003.

35. Wagner, N., Löhler, J., Kunkel, E.J., Ley, K., Leung, E., Krissansen, G., Rajewsky, K., and Müller, W. (1996). Critical role for β7 integrins in formation of the gut-associated lymphoid tissue. Nature 382, 366–370. 10.1038/382366a0.

36. Biton, M., Haber, A.L., Rogel, N., Burgin, G., Beyaz, S., Schnell, A., Ashenberg, O., Su, C.-W., Smillie, C., Shekhar, K., et al. (2018). T Helper Cell Cytokines Modulate Intestinal Stem Cell Renewal and Differentiation. Cell 175, 1307–1320.e22. 10.1016/j.cell.2018.10.008.

37. Koyama, M., Mukhopadhyay, P., Schuster, I.S., Henden, A.S., Hülsdünker, J., Varelias, A., Vetizou, M., Kuns, R.D., Robb, R.J., Zhang, P., et al. (2019). MHC Class II Antigen Presentation by the Intestinal Epithelium Initiates Graft-versus-Host Disease and Is Influenced by the Microbiota. Immunity 51, 885–898.e7. 10.1016/j.immuni.2019.08.011.

38. Bilate, A.M., London, M., Castro, T.B.R., Mesin, L., Bortolatto, J., Kongthong, S., Harnagel, A., Victora, G.D., and Mucida, D. (2020). T Cell Receptor Is Required for Differentiation, but Not Maintenance, of Intestinal CD4+ Intraepithelial Lymphocytes. Immunity 53, 1001–1014.e20. 10.1016/j.immuni.2020.09.003.

39. Brabec, T., Schwarzer, M., Kováčová, K., Dobešová, M., Schierová, D., Březina, J., Pacáková, I., Šrůtková, D., Ben-Nun, O., Goldfarb, Y., et al. (2023). Segmented filamentous bacteria–induced epithelial MHCII regulates cognate CD4+ IELs and epithelial turnover. J. Exp. Med. 221, e20230194. 10.1084/jem.20230194.

40. Suzuki, S., Miyamoto, K., Tojo, A., Yoshimatsu, Y., Teratani, T., Uchida, H., Nemoto, Y., Okamoto, R., Sihombing, A.M., Sato, T., et al. (2026). Intestinal epithelial MHC class II induces encephalitogenic CD4 T cells and initiates central nervous system autoimmunity. Sci. Immunol. 11, eaec1627. 10.1126/sciimmunol.aec1627.

41. Holiar, V., Rudenko, V., Weller, C., Naumova, M., Lebon, S., Canella, M., Busko, P., Sarusi-Portuguez, A., Shalit, T., Menachem, A.H., et al. (2026). Epithelial MHC II antigen presentation dynamically informs intestinal homeostasis and injury. bioRxiv, 2026.03.18.712348. 10.64898/2026.03.18.712348.

42. Lockhart, A., Reed, A., Castro, T.R. de, Herman, C., Canesso, M.C.C., and Mucida, D. (2023). Dietary protein shapes the profile and repertoire of intestinal CD4+ T cells. J. Exp. Med. 220, e20221816. 10.1084/jem.20221816.

43. Ansaldo, E., Yong, D., Carrillo, N., McFadden, T., Abid, M., Corral, D., Rivera, C., Farley, T., Bouladoux, N., Gribonika, I., et al. (2026). T-bet-expressing Tr1 cells driven by dietary signals dominate the small intestinal immune landscape. Proc. Natl. Acad. Sci. 123, e2520747122. 10.1073/pnas.2520747122.

44. Guehler, S.R., Finch, R.J., Bluestone, J.A., and Barrett, T.A. (1998). Increased threshold for TCR-mediated signaling controls self reactivity of intraepithelial lymphocytes. J. Immunol. (Baltim., Md : 1950) 160, 5341–5346.

45. Cheroutre, H., Lambolez, F., and Mucida, D. (2011). The light and dark sides of intestinal intraepithelial lymphocytes. Nat. Rev. Immunol. 11, 445–456. 10.1038/nri3007.

46. Bunker, J.J., Blum, J.E., Meng, X., Lopez, E.M., Weakley, A.M., Cabrera, A.V., Higginbottom, S., Kong, R., Schulman, E.A., Sattely, E.S., et al. (2026). Microbiota- and diet-specific T cells become Tregs by default. bioRxiv, 2026.08.25.747099. 10.64898/2026.08.25.747099.

47. Vetter, L., Witchley, J.N., Ahn, S.H., Brewer, R.C., Ly, A., Tout, A., Wagner, A., and Barton, G.M. (2026). Profiling of intestinal T cells reveals antigen-specific dependency on early-life programming for the generation of microbiota-reactive Tregs. bioRxiv, 2026.09.23.753882. 10.64898/2026.09.23.753882.

48. Hadis, U., Wahl, B., Schulz, O., Hardtke-Wolenski, M., Schippers, A., Wagner, N., Müller, W., Sparwasser, T., Förster, R., and Pabst, O. (2011). Intestinal Tolerance Requires Gut Homing and Expansion of FoxP3+ Regulatory T Cells in the Lamina Propria. Immunity 34, 237–246. 10.1016/j.immuni.2011.01.016.

49. Kim, K.S., Hong, S.-W., Han, D., Yi, J., Jung, J., Yang, B.-G., Lee, J.Y., Lee, M., and Surh, C.D. (2016). Dietary antigens limit mucosal immunity by inducing regulatory T cells in the small intestine. Science 351, 858–863. 10.1126/science.aac5560.

50. Blum, J.E., Kong, R., Schulman, E.A., Chen, F.M., Upadhyay, R., Romero-Meza, G., Littman, D.R., Fischbach, M.A., Nagashima, K., and Sattely, E.S. (2026). Identification and characterization of dietary antigens in oral tolerance. Sci. Immunol. 11, eaeb4684–eaeb4684. 10.1126/sciimmunol.aeb4684.

51. Chi, X., Wang, C.H., Parisotto, Y.F., Nyberg, W.A., Cabric, V., Gelineau, A., Cao, Y., Owen, D.L., Ambjörnsson, J., Mathis, D., et al. (2026). TCR origin and specificity to environmental or self-antigens on distinct antigen-presenting cells determine peripheral Treg cell differentiation. Immunity 59, 2266–2281.e6. 10.1016/j.immuni.2026.07.001.

52. Schindelin, J., Arganda-Carreras, I., Frise, E., Kaynig, V., Longair, M., Pietzsch, T., Preibisch, S., Rueden, C., Saalfeld, S., Schmid, B., et al. (2012). Fiji: an open-source platform for biological-image analysis. Nat. Methods 9, 676–682. 10.1038/nmeth.2019.

53. Gur-Cohen, S., Yang, H., Baksh, S.C., Miao, Y., Levorse, J., Kataru, R.P., Liu, X., Cruz-Racelis, J. de la, Mehrara, B.J., and Fuchs, E. (2019). Stem cell–driven lymphatic remodeling coordinates tissue regeneration. Sci. (N. York, NY) 366, 1218–1225. 10.1126/science.aay4509.

54. Mach, N., Gillessen, S., Wilson, S.B., Sheehan, C., Mihm, M., and Dranoff, G. (2000). Differences in dendritic cells stimulated in vivo by tumors engineered to secrete granulocyte-macrophage colony-stimulating factor or Flt3-ligand. Cancer Res. 60, 3239–3246.

55. Brochet, X., Lefranc, M.-P., and Giudicelli, V. (2008). IMGT/V-QUEST: the highly customized and integrated system for IG and TR standardized V-J and V-D-J sequence analysis. Nucleic Acids Res. 36, W503–W508. 10.1093/nar/gkn316.

56. Fleming, S.J., Chaffin, M.D., Arduini, A., Akkad, A.-D., Banks, E., Marioni, J.C., Philippakis, A.A., Ellinor, P.T., and Babadi, M. (2023). Unsupervised removal of systematic background noise from droplet-based single-cell experiments using CellBender. Nat. Methods 20, 1323–1335. 10.1038/s41592-023-01943-7.

57. Hao, Y., Stuart, T., Kowalski, M.H., Choudhary, S., Hoffman, P., Hartman, A., Srivastava, A., Molla, G., Madad, S., Fernandez-Granda, C., et al. (2024). Dictionary learning for integrative, multimodal and scalable single-cell analysis. Nat. Biotechnol. 42, 293–304. 10.1038/s41587-023-01767-y.

58. Wolock, S.L., Lopez, R., and Klein, A.M. (2019). Scrublet: Computational Identification of Cell Doublets in Single-Cell Transcriptomic Data. Cell Syst. 8, 281–291.e9. 10.1016/j.cels.2018.11.005.

59. Wickham, H. (2016). ggplot2, Elegant Graphics for Data Analysis. 10.1007/978-3-319-24277-4.

60. Marsh, S., Hoffman, P., kew24, and Pir, M.S. (2026). scCustomize: Release 3.3.0 (GitHub).

61. Schöneich, J., Dupont, A., Schlößer, S., Cheng, M., Schmitz, M.A., Richter, I., Gubbi, N.M.K.P., Zhang, K., Maié, T., Laouina, A., et al. (2026). Developmental factors drive the compartmentalized and discontinuous maturation of the small intestinal epithelium during the early postnatal period. PLOS Biol. 24, e3003212. 10.1371/journal.pbio.3003212.

62. Chu, T., Wang, Z., Pe’er, D., and Danko, C.G. (2022). Cell type and gene expression deconvolution with BayesPrism enables Bayesian integrative analysis across bulk and single-cell RNA sequencing in oncology. Nat. Cancer 3, 505–517. 10.1038/s43018-022-00356-3.

