## Supplementary material for "Weaning imprints a hybrid cytotoxic-regulatory identity on intestinal CD4^+^ T lymphocytes"

**SUPPLEMENTARY MATERIALS:**

Methods
Supplementary Figures 1-6
Legends for Supplementary Figures 1-6

**STAR★METHODS**

**KEY RESOURCES TABLE**

| REAGENT or RESOURCE | SOURCE | IDENTIFIER |
|---|---|---|
| Antibodies | | |
| FITC anti-human/mouse CD44 (clone: IM7) | Thermo Fisher Scientific | Cat# 11-0441-85; RRID:AB_465046 |
| PerCP-eFluor710 anti-mouse TCRγδ (clone: GL-3, GL3) | Thermo Fisher Scientific | Cat# 46-5711-82; RRID:AB_2016707 |
| APC-eFluor780 anti-mouse TCRβ (clone: H57-597) | Thermo Fisher Scientific | Cat# 47-5961-82; RRID:AB_1272173 |
| Alexa Fluor 700 anti-mouse CD45 (clone: 30-F11) | Thermo Fisher Scientific | Cat# 56-0451-82; RRID:AB_891454 |
| BV605 anti-mouse CD62L (clone: MEL-14) | BioLegend | Cat# 104438; RRID:AB_2563058 |
| BV711 anti-mouse CD8β (clone: H35-17.2) | BD Biosciences | Cat# 740761; RRID:AB_2740424 |
| BV785 anti-mouse CD103 (clone: 2E7) | BioLegend | Cat# 121439; RRID:AB_2800588 |
| BV805 anti-mouse CD8α (clone: 53-6.7) | BD Biosciences | Cat# 612898; RRID:AB_2870186 |
| BUV496 anti-mouse CD4 (clone: GK1.5) | BD Biosciences | Cat# 612952; RRID:AB_2813886 |
| APC anti-human Granzyme B (clone: GB11) | Thermo Fisher Scientific | Cat# GRB05; RRID:AB_2536539 |
| Alexa Fluor 647 anti-mouse RORγt (clone: Q31-378) | BD Biosciences | Cat# 562682; RRID:AB_2687546 |
| eFluor 450 anti-mouse/rat Foxp3 (clone: FJK-16 s) | Thermo Fisher Scientific | Cat# 48-5773-82; RRID:AB_1518812 |
| BV421 anti-mouse/rat RORγt (clone: Q31-378) | BD Biosciences | Cat# 562894; RRID:AB_2687545 |
| BV421 anti-mouse IL-17A (clone: TC11-18H10.1) | BioLegend | Cat# 506925; RRID:AB_10900442 |
| BUV395 anti-mouse/rat Ki-67 (clone: SolA15) | Thermo Fisher Scientific | Cat# 363-5698-80; RRID:AB_2925288 |
| PE-Cy7 anti-human/mouse T-bet (clone: 4B10) | Thermo Fisher Scientific | Cat# 25-5825-80; RRID:AB_11041809 |
| PE-Cy7 anti-mouse IFNγ (clone: XMG1.2) | Thermo Fisher Scientific | Cat# 25-7311-82; RRID:AB_469680 |
| APC-CY7 anti-mouse CD45 (clone: 30-F11) | BioLegend | Cat# 103116; RRID:AB_312981 |
| eFluor 660 anti-mouse LYVE1(clone: ALY7) | Thermo Fisher Scientific | Cat# 50-0443-82; RRID:AB_10597449 |
| PE anti-mouse PDPN (clone: 8.1.1) | BioLegend | Cat# 127408; RRID:AB_2161928 |

| | | |
|---|---|---|
| PerCP-Cy5.5 anti-mouse PDPN (clone: 8.1.1) | BioLegend | Cat# 127422; RRID:AB_2814016 |
| PE-Cy7 anti-mouse CD326 (clone: G8.8) | BioLegend | Cat# 118216; RRID:AB_1236471 |
| APC anti-mouse CD326 (clone: G8.8) | BioLegend | Cat# 118214; RRID:AB_1134102 |
| PE-Cy7 anti-mouse CD31 (clone: MEC13.3) | BioLegend | Cat# 102524; RRID:AB_2572182 |
| PE-Cy7 anti-mouse CD31 (clone: MEC13.3) | BioLegend | Cat# 102506; RRID: AB_312913 |
| BV421 anti-mouse/human CD11b (clone: M1/70) | BioLegend | Cat# 101235; RRID:AB_10897942 |
| PE-Cy7 anti-mouse CD11c (clone: N418) | BioLegend | Cat# 117318; RRID:AB_493568 |
| PE anti-mouse Ly-6G (clone: 1A8) | BioLegend | Cat# 127608; RRID:AB_1186099 |
| FITC anti-mouse Ly-6C (clone: HK1.4) | BioLegend | Cat# 128006; RRID:AB_1186135 |
| BV605 anti-mouse CD64 (clone: X54-5/7.1) | BioLegend | Cat# 139323; RRID:AB_2629778 |
| Alexa Fluor 700 anti-mouse I-A/I-E (clone: M5/114.15.2) | BioLegend | Cat# 107622; RRID:AB_493727 |
| BV605 anti-mouse I-A/I-E (clone: M5/114.15.2) | BioLegend | Cat# 107639; RRID:AB_2565894 |
| Alexa Fluor 488 anti-mouse TCR γ/δ (clone: GL3) | BioLegend | Cat# 118128; RRID:AB_2562771 |
| PerCP-Cy5.5 anti-mouse CD90.2 (clone: 30-H12) | BioLegend | Cat# 105338; RRID:AB_2571945 |
| Alexa Fluor 488 anti-mouse CD90.2 (clone: 30-F11) | BioLegend | Cat# 105316; RID:AB_492886 |
| PE anti-mouse CD8α (clone: 53-6.7) | BioLegend | Cat# 100707; RRID:AB_312746 |
| PE-Cy7 anti-mouse CD4 (clone: GK1.5) | BioLegend | Cat# 100422; RRID:AB_312707 |
| BV421 anti-mouse CD3ε (clone: 145-2C11) | BioLegend | Cat# 100335; RRID:AB_10898314 |
| APC anti-mouse Itgα4β7 (clone: DATK32) | BioLegend | Cat# 120607; RRID:AB_10719833 |
| PE-Cy7 anti-mouse CD4 (clone: GK1.5) | Thermo Fisher Scientific | Cat# 25-0041-81; RRID:AB_469575 |
| PE anti-mouse CD45.1 (clone: A20) | Thermo Fisher Scientific | Cat# 12-0453-83; RRID:AB_465676 |
| APC anti-mouse TCRβ (clone: H57-597) | Thermo Fisher Scientific | Cat# 17-5961-83; RRID:AB_469482 |
| E-Cadherin Rabbit mAb (clone: 24E10) | Cell Signaling Technology | Cat# 3195S; RRID:AB_2291471 |

| | | |
|---|---|---|
| Alexa Fluor 488 anti-mouse CD45 (clone: 30-F11) | BioLegend | Cat# 103122; RRID:AB_493531 |
| Anti-PECAM-1 Antibody, Azide Free (clone: 2H8) | Millipore | Cat# MAB1398Z; RRID: AB_94207 |
| Alexa Fluor 647 anti-mouse CD3 (clone: 17A2) | BioLegend | Cat# 100209; RRID:AB_389323 |
| CD3e Monoclonal Antibody (clone: 145-2C11) | Thermo Fisher Scientific | Cat# 14-0031-82; RRID:AB_467049 |
| PE anti-mouse CD4 (clone RM4-5) | BioLegend | Cat# 100512; RRID:AB_312715 |
| Alexa Fluor 561 anti-CD4 (clone RM4-5) | Thermo Fisher Scientific | Cat# 505-0042-82; RRID:AB_2896185 |
| Alexa Fluor 647 anti-mouse CD8α (clone: 53.6-7) | BioLegend | Cat# 100724; RRID:AB_389326 |
| Alexa Fluor 488 anti-mouse TCR γ/δ (clone: GL3) | BioLegend | Cat# 118128; RRID: AB_2562771 |
| Alexa Fluor 488 anti-mouse CD206 (clone: C068C2) | BioLegend | Cat# 141710; RRID:AB_10900445 |
| Alexa Fluor 647 anti-mouse Tim-4 (clone: RMT4-54) | BioLegend | Cat# 130008; RRID:AB_2271648 |
| eFluor 660 anti-CD324 (E-Cadherin) (clone: DECMA-1) | Thermo Fisher Scientific | Cat# 50-3249-82; RRID:AB_11040003 |
| Alexa Fluor 647 anti-mouse I-A/I-E (clone: M5/114.15.2) | BioLegend | Cat# 107618; RRID:AB_493525 |
| Mouse MAdCAM-1 Antibody | R&D Systems | Cat# AF993; RRID:AB_355772 |
| Rabbit anti-RFP | Elaine Fuchs's lab | NA |
| Alexa Fluor 647 anti-human/mouse Granzyme B (clone: GB11) | BioLegend | Cat# 372220; RRID:AB_2728387 |
| Alexa Fluor 647 Goat anti-hamster (Armenian) IgG (clone: Poly4055) | BioLegend | Cat# 405510; RRID:AB_2566695 |
| Rhodamine Red-X (RRX) AffiniPure Donkey Anti-Rabbit IgG (H+L) | Jackson ImmunoResearch | Cat# 711-295-152; RRID:AB_2340613 |
| Rhodamine Red-X (RRX) AffiniPure Donkey Anti-Goat IgG (H+L) | Jackson ImmunoResearch | Cat# 705-295-003; RRID: AB_2340422 |
| Anti-mouse MHC-II (I-A/I-E) | Bio X Cell | Cat# BE0108; RRID:AB_10949298 |
| TruStain FcX (anti-mouse CD16/32) Antibody | Biolegend | Cat# 101320; RRID:AB_1574975 |
| Bacterial and virus strains | | |
| Selected bacterial strains organized as per their timing of intestinal colonization – Table S6 | NA | NA |
| All bacterial strains (and other antigens) tested for TCR reactivity – Table S7 | NA | NA |
| Biological samples | | |

| | | |
|---|---|---|
| Fetal Bovine Serum | Sigma-Aldrich | F0926 |
| Newborn Calf Serum | Thermo Fisher Scientific | 16010-159 |
| Chemicals, peptides, and recombinant proteins | | |
| EDTA, 0.5M, pH8.0 | Thermo Fisher Scientific | AM9260G |
| Percoll Centrifugation Media | Cytiva | 17089101 |
| Corn oil | Sigma-Aldrich | C8267 |
| Dithiothreitol (DTT) | Thermo Fisher Scientific | R0862 |
| Collagenase VIII | Sigma-Aldrich | C-2139 |
| Collagenase D | Roche | 11088882001 |
| DNAse I | Roche | 10104159001 |
| HEPES 1M | Thermo Fisher Scientific | 15630080 |
| MEM NEAA (100X) | Thermo Fisher Scientific | 11140-050 |
| Sodium pyruvate (100 mM) | Thermo Fisher Scientific | 11360-070 |
| RPMI 1640 | Thermo Fisher Scientific | 21870076 |
| 2-Mercaptoethanol (BME) | Sigma-Aldrich | M6250 |
| Phorbol Myristate Acetate | Sigma-Aldrich | P8139 |
| Ionomycin Calcium Salt from Streptomyces conglobatus | Sigma-Aldrich | IO634 |
| BD GolgiStop Protein Transport Inhibitor (containing Monensin) | BD Biosciences | 554724 |
| Tamoxifen | Sigma-Aldrich | T5648-5G |
| DMEM (1X) | Thermo Fisher Scientific | 11960-044 |
| Penicillin-Streptomycin (Pen Strep) | Thermo Fisher Scientific | 15140-122 |
| L-Glutamine | Thermo Fisher Scientific | 25030-081 |
| Pen Strep Glutamine (100X) | Thermo Fisher Scientific | 10378-016 |
| Protamine Sulfate | MP Biomedicals | 194729 |
| Puromycin | Thermo Fisher Scientific | A11138-03 |
| Blasticidin S HCl | Thermo Fisher Scientific | A11139-03 |
| IMDM (1X) + GlutaMAX-I | Thermo Fisher Scientific | 31980-030 |
| TrypLE Express | Thermo Fisher Scientific | 12605-010 |
| Red Blood Cell Lysing Buffer Hybri-Max™ | Sigma-Aldrich | R7757 |

| | | |
|---|---|---|
| Critical commercial assays | | |
| LIVE/DEAD Fixable Aqua Dead Cell Stain | Thermo Fisher Scientific | L34966 |
| Zombie NIR Fixable Viability Kit | BioLegend | 423106 |
| eBioscience™ Foxp3 / Transcription Factor Staining Buffer Set | Thermo Fisher Scientific | 00-5523-00 |
| BD Cytofix/Cytoperm | BD Biosciences | 51-2090KZ |
| PBS57-loaded CD1d tetramer | NIH Tetramer Core Facility | IEDB ID: 182375 |
| Lipofectamine 3000 | Thermo Fisher Scientific | L3000001 |
| CD11c MicroBeads | Miltenyi-Biotec | 130-125-835 |
| Chromium Next GEM Single Cell 5' HT Kit v2 | 10X Genomics | 1000374 |
| Deposited data | | |
| scRNA-seq analysis | This paper | GEO: GSE[XXXXXX] |
| Raw single and spatial transcriptomic data (mouse small and large intestine) | Niec et al.[19] | GEO: GSE190037 |
| Experimental models: Cell lines | | |
| NFAT-GFP 58$\alpha^{-}\beta^{-}$ hybridoma | K. Murphy | NA |
| Platinum-E (Plat-E) Retroviral Packaging Cell Line | Cell Biolabs | Cat# RV-101, RRID:CVCL_B488 |
| B16-Flt3L melanoma cells | M. Nussenzweig | NA |
| Experimental models: Organisms/strains | | |
| Mouse: C57BL/6J | Jackson Laboratory | JAX: 000664 |
| Mouse: C57BL/6J-*Ptprc*$^{em6Lutzy}$/J | Jackson Laboratory | JAX: 033076 |
| Mouse: B6.129P2-*Lgr5*$^{tm1(cre/ERT2)Cle}$/J | Jackson Laboratory | JAX: 008875 |
| Mouse: B6.129S7-*Rag1*$^{tm1Mom}$/J | Jackson Laboratory | JAX: 002216 |
| Mouse: B6.Cg-*Rag2*$^{tm1.1Cgn}$/J | Jackson Laboratory | JAX: 008449 |
| Mouse: B6.129(Cg)-*Gt(ROSA)26Sor*$^{tm4(ACTB-tdTomato,-EGFP)Luo}$/J | Jackson Laboratory | JAX: 007676 |
| Mouse: B6.Cg-*Gt(ROSA)26Sor*$^{tm14(CAG-tdTomato)Hze}$/J | Jackson Laboratory | JAX: 007914 |
| Mouse: *Sell*$^{Cre-ERT2}$ | M. Nussenzweig, The Rockefeller University | NA |
| Oligonucleotides | | |
| Primer: *Ccl25* forward: 5'- CCGGCATGCTAGGAATTATCA – 3' | Integrated DNA Technologies | NA |

| | | |
|---|---|---|
| Primer: *Ccl25* reverse:<br>5'- GGCACTCCTCACGCTTGTACT – 3' | Integrated DNA Technologies | NA |
| Primer: *H2-2a* forward:<br>5'- TCAGTCGCAGACGGTGTTTAT – 3' | Integrated DNA Technologies | NA |
| Primer: *H2-2a* reverse:<br>5'- GGGGGCTGGAATCTCAGGT – 3' | Integrated DNA Technologies | NA |
| Primer: *H2-Ab1* forward:<br>5'- AGCCCCATCACTGTGGAGT – 3' | Integrated DNA Technologies | NA |
| Primer: *H2-Ab1* reverse:<br>5'- GATGCCGCTCAACATCTTGC – 3' | Integrated DNA Technologies | NA |
| Primers used for scTCR-seq – Table S8 | Integrated DNA Technologies | NA |
| Recombinant DNA | | |
| pMSCV-mCD4-PIG TCR-OTII | Nagashima et al.[10] | NA |
| TCR clonotype constructs – Table S4 | | |
| Software and algorithms | | |
| FlowJo v10+ | TreeStar | https://www.flowjo.com/ |
| GraphPad Prism v9+ | GraphPad | https://www.graphpad.com/ |
| Imaris 9.5 | Oxford Instruments | https://imaris.oxinst.com/versions/9-5 |
| Fiji | Schindelin et al.[52] | https://imagej.net/software/fiji/downloads |
| BioRender | BioRender | https://www.biorender.com/ |
| Claude | Anthropic | https://claude.com/ |
| Claude Code | Anthropic | https://claude.com/product/claude-code |
| Claude Science | Anthropic | https://claude.com/product/claude-science |
| Other | | |
| 27 G x 1/2 in. BD PrecisionGlide™ Needle | BD | 305109 |
| Plastic feeding tubes, 22ga (black) x 25mm | Instech Laboratories | FTP-22-25-50 |
| Plastic feeding tubes, 20ga (yellow) x 38mm | Instech Laboratories | FTP-20-38 |

#### EXPERIMENTAL MODEL DETAILS

##### Mouse models

All experiments were performed in specific-pathogen-free (SPF) or germ-free (GF) mice on the C57BL/6 background. SPF C57BL/6J (Jax 000664), CD45.1 ($Ptprc^{K302E}$; Jax 033076), *Lgr5-EGFP-IRES-Cre*$^{ERT2}$ (Jax 008875), *Rag1*$^{-/-}$ (Jax 002216), *Rag2*$^{-/-}$ (Jax 008449), B6.129(Cg)-*Gt(ROSA)26Sor*$^{tm4(ACTB\text{-}tdTomato,\text{-}EGFP)Luo}$/J (Jax 007676), and *Rosa26*$^{CAG\text{-}LSL\text{-}tdTomato}$ (Ai14, Jax 007914) mice were purchased from Jackson Laboratories. Ai14 mice were interbred with *Sell*$^{Cre\text{-}ERT2}$ mice, provided by M. Nussenzweig[23], to generate Sell-Tomato mice for timestamping and fate-mapping analyses. Both male and female mice were used for all experiments. Mice were maintained in the Association for Assessment and Accreditation of Laboratory Animal Care-accredited animal facility of The Rockefeller University (RU) under SPF or GF conditions, and procedures were performed with Institutional Animal Care and Use Committee approved protocols (IACUC protocol 25036-H). GF animals were housed in the Mucida Laboratory germ-free facility at The Rockefeller University. GF mice were exported and housed in ISOcages (Tecniplast Inc.) for experiments. Pet store mice were housed in a separate isolator/room at the Rockefeller animal facility. Mice of all strains were housed in an environment with controlled temperature and humidity, on a 12-hour light cycle, and fed regular chow, unless noted otherwise. The numbers of animals used for each experiment are specified in the figure legends per timepoint analyzed. Adult animals were used at 3-13 weeks of age for experiments, while postnatal ages of neonates are indicated by day in the respective figure or figure legends. Animals were co-housed and littermate controls were used, whenever applicable.

#### METHOD DETAILS

##### Tamoxifen treatments

For timestamping and fate-mapping of naïve T cells in Sell-Tomato mice (SPF and GF), tamoxifen (Sigma-Aldrich) was dissolved in sterile corn oil (Sigma-Aldrich) and filtered in 0.22µm filter prior to administration to mice. Unless otherwise specified, SPF neonates received a pulse of tamoxifen split into two doses two days apart (1 mg/mL, 50 µL/mouse) through intragastric milk-spot injection using a 27G x ½ needle (BD) at postnatal days 1 (P1) and 3 (P3). P6-P12 SPF mice were administered tamoxifen (1 mg/mL, 100 µL/mouse) via gavage. P13-P21 mice received 50 µL of 20 mg/mL solution; P22-6 week-old mice received 100 µL of 20 mg/mL solution, whereas mice older than 6 weeks of age received 100 µL of 50 mg/mL solution. In all cases, mice were orally administered tamoxifen via gavage (feeding tubes Instech Laboratories). GF neonates, weanlings, and adults were administered tamoxifen in a single dose (double volume) to minimize risk of potential contamination during manipulation.

##### Isolation of T cells from the intestine and mesenteric lymph nodes

Intraepithelial (IE) and lamina propria (LP) lymphocytes from weanling and adult mice were isolated as previously described[38]. Briefly, small intestines were harvested and washed in PBS, then incubated in PBS+1mM dithiothreitol (DTT, Thermo Fisher Scientific), followed by two washes with 30 mM EDTA (Thermo Fisher Scientific). IE cells were recovered from the supernatant of DTT and the first EDTA wash. Lymphocytes from the lamina propria were obtained after collagenase VIII (2 mg/mL, Sigma-Aldrich) digestion of the tissue. Mononuclear cells were isolated by density gradient centrifugation using Percoll. Isolation of neonatal (up to P17) intestinal lymphocytes was modified as follows: only one EDTA wash was performed following the DTT wash, and the supernatant from washes was filtered through 70 μm cell strainers instead of metal strainers. The tissue was digested with collagenase and incubated at 80 rpm and 37°C for 25 min instead of 45 min. Mesenteric lymph nodes were harvested, cleaned of surrounding fat and smashed through 70 μm cell strainers. Single-cell suspensions were then stained with fluorescently labeled antibodies for 25 min at 4°C prior to downstream flow cytometry (analysis or sorting) as specified in figure legends.

**Staining and gating strategies**

The following gating strategies were used to analyze timestamped $CD4^+$ $Tomato^+$ T cells: single live lymphocytes (based on size and live/dead fixable dye Aqua stain), $CD45^+$, $TCR\gamma\delta^-$, TCRβ+, $CD8\beta^{-/low}$, $CD4^+$, $Tomato^+$. For analysis of timestamped/fate-mapped $CD4^+$ $Tomato^+$ T cells, the following gating strategy was used: single live lymphocytes (based on size and live/dead fixable dye Aqua stain), $CD45^+$, $TCR\gamma\delta^-$, $TCR\beta^+$, $CD8\beta^{-/low}$, $CD4^+$, $CD62L^-$, $Tomato^+$. For index-sorting of time-stamped and fate-mapped $CD4^+$ T cells for plate-based scTCR-seq, we sorted IEL samples using the following strategy: single live lymphocytes (based on size and live/dead fixable dye Aqua stain), $CD45^+$, $TCR\gamma\delta^-$, $TCR\beta^+$, $CD8\beta^{-/low}$, $CD4^+$, $CD62L^-$, $Tomato^+$, while LPL samples were sorted using the following approach: single live lymphocytes (based on size and live/dead fixable dye Aqua stain), $CD45^+$, $TCR\gamma\delta^-$, $TCR\beta^+$, $CD8\beta^{-/low}$, $CD4^+$, $CD62L^-$, $Tomato^+$, $CD8\alpha^-$ (to exclude potential IEL contaminants in the LP preparation). For 10X scRNA-seq coupled with scTCR-seq, we used the following approach: single live lymphocytes (based on size and live/dead fixable dye Aqua stain), $TCR\gamma\delta^-$, $TCR\beta^+$, CD1d $tetramer^-$ (loaded with PBS57, provided by the NIH Tetramer Core Facility), $CD4^+$, $CD62L^{-/low}$, $Tomato^+$.

**Intranuclear and Intracellular Staining and Flow Cytometry**

Intranuclear staining for Foxp3, RORγt, T-bet, and Ki-67 was performed using the Foxp3 /Transcription Factor Staining Buffer Set according to the manufacturer's instructions (eBioscience, USA). For analysis of cytokine secretion, total mononuclear cells isolated from the epithelium and lamina propria were plated in 48-well plates and incubated at 37°C with 100 ng/mL phorbol 12-myristate 13-acetate (PMA, Sigma-Aldrich) and 200 ng/mL ionomycin (Sigma-Aldrich) for 4 h. Golgi stop solution containing Monensin (2 mM, BD Biosciences) was added 1h after PMA and ionomycin. Intracellular staining for IFNγ and granzyme B was conducted in Perm/Wash buffer after fixation and permeabilization in Fix/Perm buffer (BD Biosciences, USA) according to kit instructions.

**Flow cytometry**

Fluorescence-activated cell sorting (FACS) Cell sorting for all experiments was performed at the Rockefeller Flow Cytometry Resource Centers on a BD FACSAria/Symphony S5 Cell Sorter (equipped with Diva Software v.8.0) or in-house with a Sony Biotechnology MA800. All flow cytometry experiments were run on an LSR Fortessa Analyzer (BD Biosciences), LSR-II flow cytometer (BD Biosciences), or FACSymphony A5 Cell Analyzer (BD Biosciences). Data were exported as FCS 3.0 files and analyzed in FlowJoTM v.10 software.

**Adoptive transfers**

For adoptive transfer experiments, we transferred $Tomato^+$ T cells from the mesenteric lymph nodes and spleen of neonatal (P7) or adult (>7 weeks of age) *mT/mG* or Sell-Tomato mice into neonatal (P7) or adult (>7 weeks of age) into *Rag1*$^{-/-}$, *Rag2*$^{-/-}$, or congenic CD45.1 recipients. $5x10^5$-$10^6$ Tomato$^+$ cells, resuspended in sterile PBS, were intravenously injected into neonate or adult recipients. The same number of T cells were transferred across recipients per experiment. Animals were sacrificed for tissue harvesting and imaging 1 week after the transfer.

**Immunofluorescence staining on tissue sections**

For immunofluorescence and confocal analysis, mouse intestines were dissected, flushed with ice-cold PBS and fixed with BD Cytofix/CytoPerm (Thermo Fisher Scientific) diluted 1:4 in PBS (1% PFA final concentration) at 4°C overnight. Next, the tissue was washed three times with PBS and incubated with 30% sucrose overnight at 4ºC. The tissue was embedded in OCT (Tissue Tek) the following day. Frozen tissue blocks were sectioned at 18-25 µm on a cryostat (Leica CM3050S) and mounted on SuperFrost Plus slides (Fisher). The tissue sections were blocked for 30-60 minutes at room temperature in blocking solution (PBST with 2.5% normal donkey serum [NDS], 1% BSA, 2.5% gelatin) or BD Cytoperm buffer (1:10 diluted in PBS, Thermo Fisher Scientific) with TruStain FcX (anti-mouse CD16/32) block (1:1000) (BioLegend). Sections were incubated with primary antibodies diluted in blocking solution at 4°C overnight or 1 hour at RT. Sections were washed three times with PBS and, if non-conjugated primary antibodies were used, incubated with secondary antibodies conjugated with Alexa-488, Rhodamine Red-X (RRX) or Alexa-647 (1:800, Life Technologies) in blocking solution at room temperature for 2 hours. Finally, sections were washed three times with PBS and mounted with ProLong Gold, Diamond Anti-fade Mountant (Thermo Fisher Scientific) or Fluoromount G (ThermoFisher Scientific).

**Iterative bleaching extends multiplexity (IBEX) staining**

Iterative bleaching extends multiplexity immunofluorescence stainings were performed as previously described[18]. Briefly, tissues were fixed in 1% BD Cytofix/CytoPerm (BD Biosciences) at 4°C overnight, before being switched to a 30% sucrose solution for at least 16h and embedded in OCT (Tissue Tek). Two-chamber glass slides were coated with chrome gelatin alum (Newcomersupply, cat# 1033A) at 37°C overnight and equilibrated to cryostat temperature. 25 µm sections were placed on slides, dried for 1h at 37°C, re-hydrated in PBS for 5 minutes at RT, blocked for 10 minutes in PBS 0.1% Triton followed by 30 minutes in BD Cytoperm buffer (1:10 diluted in PBS, Thermo Fisher Scientific) with TruStain FcX (anti-mouse CD16/32) block (1:1000) (BioLegend, cat#101320). Antibody mixes were prepared in said BD Cytoperm buffer with FcX block and applied for 1h at 37°C or overnight at 4°C. Slides were washed 3x in PBD, incubated with PBS + DAPI (1:1000 of a 1 mg/ml stock) for 15 minutes at RT, washed again and mounted in Fluoromount G (Thermo Fisher Scientific) to image samples on a Dragonfly spinning disc confocal microscope. After imaging, mounting media was replaced by 5 PBS washes. For iterative bleaching rounds, 1 mg/ml of lithium borohydride (prepared fresh every 4h) was used at RT for 15 minutes, followed by 3 washes in PBS and the next staining round. IBEX image tiles were stitched using the Andor Fusion software (v2.3) and later aligned using DAPI as a fiducial frame in Imaris and the SimpleITK Image Registration Pipeline plug-in for Imaris. If necessary, the channels were manually further aligned in Imaris.

**Whole-mount immunofluorescence staining**

Dissected small intestines were flushed with ice-cold PBS. Tissues were opened longitudinally and spread with the epithelial (luminal) side up on Whatman filter paper or left as short tubes. Mouse tissues were fixed in 1% BD Cytofix/CytoPerm (BD Biosciences) at 4°C overnight, followed by an overnight fixation in Dent's solution (20% dimethyl sulfoxide (DMSO) and 80% methanol) at 4°C. Subsequently, tissue pieces were washed three times in PBS and permeabilized for at least 3 hours in PBST (PBS with 0.3% Triton (Sigma)) followed by at least 1 hour in blocking solution (PBST with 2.5% NDS, 1% BSA, 2.5% gelatin). For immunolabeling, samples were incubated with primary antibodies in PBST for 24-48 hours at room temperature followed by 8-10 30-minute washes in PBST. Samples were incubated overnight at room temperature with secondary antibodies conjugated with Alexa-488, RRX, or Alexa-647 (1:200 Life Technologies) in blocking solution with DAPI (2 µg/ml, 1:500). Samples were washed at least 8 times with PBST containing DAPI (1:500) for 4 hours at room temperature prior to tissue clearing.

**Tissue clearing**

Ethyl-cinnamate-based tissue clearing was performed as previously described with some modifications[53]. Stained tissues were transferred through 30%, 50% and 70% ethanol solutions diluted in UltraPure water and adjusted to a pH of 9.0 for 1 hour each, all on a rocker at room temperature (RT). Dehydrated tissues were then incubated twice in 100% ethanol for at least 1 hour each at RT. Tissues were next transferred into 500 µl of ethyl cinnamate (Sigma) in Eppendorf tubes (Polypropylene) for clearing overnight. Cleared tissue was mounted with ethyl cinnamate drops between 2 glass cover slips size 22x40 mm, #0 (Electron Microscopy Science), and placed in the microscope slide holder for imaging.

**Imaging, epifluorescence and confocal microscopy**

Tissue sections were imaged with the 10x or 20x/0.8 air objectives of a Leica Thunder Imager Live Cell with AFC to acquire whole-tissue (swiss roll) tiles in four channels (405, 488, 561 and 625). For images with high background, the computational clearing algorithm implemented in the Leica software was applied. The same settings were applied across all images per experiment. Cleared whole-mount tissue specimens as well as IBEX samples were imaged using the 20x air or 40x oil objective of an inverted Dragonfly 202 spinning disk confocal system (Andor Technology Inc) equipped with a 40 µm pinhole and a Leica DMi8 camera with AFC (20x air objective). Four laser lines (405, 488, 561 and 625 nm) were used for near simultaneous excitation of DAPI, Alexa-448, RRX and Alexa-647 fluorophores. The system was driven by the Andor Fusion software.

**Multi-photon microscopy of tissue explants**

Terminal ilea from neonate or adult recipient mice were flushed with ice-cold PBS and fixed in 4% PFA for 2.5 h at 4ºC. Tissues were washed and stored in PBS until imaging. On the day of imaging, ~2 mm of the tissue was cut transversally under a dissection microscope, then placed in PBS between two glass coverslips held together with vacuum grease. This tissue set up was kept on a cold metal block throughout the duration of imaging. Multiphoton imaging was performed using an Olympus FV1000 upright microscope fitted with a 25X 1.05NA Plan water-immersion objective and a Mai-Tai DeepSee Ti-Sapphire laser (Spectraphysics). tdTomato-expressing cells and second harmonic generation of tissue surrounding fibrillar collagen were imaged at $\lambda = 930$ nm. To quantify the relative z-position of $Tomato^+$ $CD3^+$ or $CD4^+$ T cells in whole-mount tissue samples or explants subjected to two-photon imaging, spots were created in Imaris (Oxford Instruments) in the respective channels (based on the mean fluorescence intensity signal).

**Image processing and analysis**

Processing and rendering of imaging data was performed on Imaris 9.5 - 10 (Bitplane) and included initial image processing, 3D-surface rendering, cell-spotting analysis as well as mean signal intensity, volumetric, distance measurements and statistical color-coding of objects. 3D surface rendering and spots analysis were based on mean signal intensity per channel and a cell diameter estimate of 7 μm. Images were acquired and processed at identical settings per experiment.

To quantify MAdCAM-1$^+$ blood vascular cells, surfaces were created on the MAdCAM-1 and PECAM-1 signal and the respective volumetric values were exported from Imaris into Excel. The MAdCAM-1 volume was plotted as % of the total PECAM-1 volume and values were averaged across images to obtain 1 value per biological replicate, as indicated. In addition, the total tissue volume was inferred from the total image volume.

To quantify the relative z-position of CD3$^+$ or CD4$^+$ T cells in whole-mount tissue samples or 2-photon images from tissue explants, spots were created in Imaris in the respective channels (based on the mean fluorescence intensity signal). Each image was manually inspected after surface creation and non-specific spots (which picked up background staining) were manually removed before further analysis. The xyz position values of individual spots were then exported into Excel and max-scaled on a per-image basis using the formula $((x-x_{min})/(x_{max}-x_{min}))$. Individual cells were plotted on a 0-1 range (y-axis) and data were pooled from biological replicates. No statistics were performed on those plots given the risk of p-value inflation due to a high number of datapoints.

These z-position plots accompany plots that show the percentage of T cells in the villus compartment in figure 1D (calculated from the same input images). For those, the number of spots (T cells) within the villus compartment of whole-mount images (crypts and villi were determined based on ECAD staining and the image was divided at the respective plane) were counted and plotted as a percentage of the total T cell number. Values were averaged per biological replicate (at least 3 images were taken per animal) and plotted accordingly. The same type of analysis was done for images from adoptive transfer experiments, but Tomato$^+$ donor cells in the recipient tissue were manually counted, given the lower number of cells. For all these analyses, the percentage values of crypt and villus add up to 100%, hence only quantification for the villus compartment are shown. The percentage of Gzmb$^+$ T cells was calculated as a fraction of the number of Gzmb$^+$ cells over total number of T cells in the crypt and villus compartment.

**Small intestinal epithelial cell sorting for rt-qPCR**

The small intestines of neonatal (P9) and adult (>7 weeks of age) SPF C57B6/J animals were dissected, flushed with ice-cold PBS and opened longitudinally. The opened intestine was gently scraped with a glass coverslip twice to remove mucus and villous epithelium, followed by additional washes of the tissue in PBS on ice. The tissue was cut into small fragments and underwent 5 additional washes in PBS by shaking. Subsequently, the tissue was incubated in 20 ml of PBS with 10 mM EDTA on a horizontal shaker at 4°C for 20 minutes, which was followed by a second incubation in PBS with 10 mM EDTA. After each incubation, the tissue was shaken vigorously and the solution was collected, filtered through a 100 μm strainer and eventually pooled. These fractions were enriched for (crypt) epithelial cells, spun down at 1300 rpm and resuspended in FACS buffer and FcX block (1:1000, Biolegend). Cells were stained for CD326-APC, MHCII-BV605, PDPN-PerCp-Cy5.5, CD45-APC-Cy7, CD24-PE and CD31-FITC (all at 1:200) and resuspended in FACS buffer with DAPI (1:30,000) before sorting. Single alive lin- (i.e.

PDPN⁻ CD45⁻ CD31⁻) CD326⁺ cells were sorted into Trizol for RNA extraction and real time (rt)-qPCR for *Ccl25*, *H2-Aa* and *H2-Ab1* genes.

**RNA isolation**
RNA was isolated from 60,000 – 100,000 small intestinal epithelial cells (lin⁻ EpCAM⁺) sorted into 300 µl of TRIzol Reagent (Thermo Fisher). RNA was extracted from cells using the Direct-zol RNA MicroPrep Kit (Zymo Research) according to manufacturer's instructions including treatment with DNase I. The final elution volume was 10 µl and RNA quantity and quality were measured on a NanoDrop 8000 spectrophotometer (Thermo Fisher). The same amount of RNA (up to 500 ng) was loaded per sample for subsequent cDNA synthesis. For RNA isolation from sorted small intestinal blood endothelial, lymphatic endothelial and fibroblast cells 100 000 cells were sorted into 300 µl of TRIzol Reagent.

**cDNA synthesis**
At least 100 ng of RNA were loaded per sample, and cDNA was synthesized using the SuperScript™ VILO™ cDNA Synthesis Kit (Thermo Fisher). Briefly, in a final volume of 20 µl we mixed VILO Reaction mix (5X), SuperScript Enzyme (10X), RNA template and DEPC-treated water (Thermo Fisher). cDNA was synthesized using the following PCR program: 10 minutes at 25°C – 60 minutes at 42°C – 5 minutes at 85°C – 12°C. For all quantitative real-time PCR (qPCR) reactions cDNA was diluted to get a final concentration of 5 ng/µl.

**Quantitative real-time PCR**
For qPCR equal amounts of cDNA template were mixed with validated gene-specific primers (amplification efficiency between 90 – 110% as tested by standard curves in advance), DEPC-treated water and Power SYBR green PCR Master Mix (Thermo Fisher). qPCR was performed in technical duplicates using an Applied Biosystems 7900HT Fast Real-Time PCR machine and relative expression (RE) to a control sample was calculated with the RelQuant software (Bio-Rad Laboratories) by normalizing to *B2m* housekeeping gene expression. Primer sequences are available under the "oligonucleotides" section in the STAR methods.

**Isolation of intestinal epithelial cells to profile levels of MHCII**
To profile the expression of MHCII by epithelial cells, the small intestine of *Lgr5-EGFP-ires-*Cre$^{ERT2}$ mice was dissected, flushed with ice-cold PBS, opened longitudinally and cut into small pieces. Tissue fragments were washed 5x in ice-cold PBS, before incubating in 20 ml PBS with 10 mM ethylenediaminetetraacetic acid (EDTA, Invitrogen) for 20 minutes on a horizontal shaker at 4°C. Tubes were vigorously shaken after this incubation, and the solution was filtered through a 100 µm strainer. The incubation was repeated; fractions were pooled and spun down at 1300 rpm. The pellets were resuspended in TrypLE Express Enzyme (1X, Gibco) solution with DNase (0.1 mg/ml, Roche) and incubated at 37°C for 1 minute under constant agitation. The cell suspension was diluted in PBS, filtered through 100 µm and 40 µm strainers, before being spun down at 1300 rpm. The pellet was resuspended in FACS buffer (PBS, 2% FBS, 1X HEPES) with TruStain FcX (anti-mouse CD16/32, BioLegend, cat#101320) block (1:1000) before proceeding with cell staining including the following markers: CD326-APC, MHCII-BV605, CD45-APC-Cy7 (all at 1:200). The cells were resuspended in FACS buffer and DAPI (1:30,000, from a 1 mg/ml stock) and run on the flow cytometer.

**Generation of T cell hybridomas**

The retroviral vector pMSCV-mCD4-PIG TCR-OTII (Twist Biosciences) was used to transduce NFAT-GFP 58α⁻β⁻ hybridoma cells with synthetic TCR constructs of interest as described previously[10,31]. Sequences for the TCRα and TCRβ genes were obtained from the filtered_contig_annotations.csv output file from scRNA-seq (Table S4). Target constructs consisting of the TCRα and TCRβ genes separated by the self-cleaving P2A linker peptide were synthesized (Twist Biosciences) and used to replace the OT-II TCR in the pMSCV-mCD4-PIG plasmid. The Platinum-E retroviral packaging cell line (RV-101, Cell Biolabs) was used for viral packaging of pMSCV-mCD4-PIG TCR vectors. Platinum-E cells were cultured in a 96-well plate with blasticidin (10 μg/mL, Thermo Fisher Scientific) and puromycin (1 μg/mL, Thermo Fisher Scientific) to 60-80% confluence. $2x10^4$ Plat-E cells were cultured in an antibiotic-free medium and transfected with pMSCV-mCD4-PIG TCR vectors as a DNA-lipid complex using Lipofectamine 3000 (L3000001, ThermoFisher Scientific). The culture supernatant containing viral particles was harvested 24 hours after transfection. Then, $2.5\times10^4$ NFAT-GFP cells were centrifuged, resuspended in 100 μl of the virus-containing supernatant supplemented with 10 μg/mL protamine sulfate (MP Biomedicals) and plated in U-bottom 96-well plates. NFAT-GFP cells were spin-transduced by centrifugation at 1,000 g at 32°C for 120 min, pipetted mixed, and cultured overnight at 37°C in a 5% $CO_2$ incubator. Cells were then transferred into a flask and incubated for 1-2 weeks with 10 mL of culture medium (DMEM with 10%FCS, penicillin–streptomycin, and 2 mM L-glutamine) under puromycin selection (1 μg/mL). After selection and propagation, TCR expression in NFAT-GFP cells was confirmed by flow cytometry.

**Antigen extract preparation**

The following classes of antigen extracts were prepared for *in vitro* screening assays: food, intestinal luminal content and feces, defined bacterial strains, and bacterial pathogens. With the exception diet-derived antigens, all antigen extracts were heat-killed at 75 °C for 1 hour on a heat block prior to use.

*Food extracts*

Food extracts were prepared from standard chow diet and its individual ingredients. Chow pellets were ground into a fine powder and suspended in PBS to make working solutions of 50 mg/mL. For individual dietary ingredients, soybean flour (Sigma-Aldrich, S9633), yeast extract (Sigma-Aldrich, Y1625), and zein were dissolved in PBS at 50 mg/mL, while gliadin was prepared at 50 mg/mL in DMSO. The dissolved dietary proteins were vortexed and stored at -20°C.

*Luminal content*

Feces and intestinal content antigens were prepared from mice at suckling, weaning, and adult ages and from three intestinal sections (duodenum, ileum, and colon). For fecal content, one small fecal pellet (<0.5 cm) was dissolved in 200 μL PBS and briefly spun on a benchtop centrifuge (5-6 s). The supernatant was moved to a clean tube and centrifuged at 10,000 x g for 10 min, and the resulting bacterial pellet was resuspended in 200 μL sterile PBS. For intestinal content, the intestine was flushed with 1 mL PBS using a gavage needle and syringe, centrifuged at 10,000 x g for 10 min, and the pellet resuspended in 200 μL PBS.

*Defined bacterial strains*

Individual bacterial antigens comprised the Oligo-MM$^{12}$ [11,32], hCom2[10] and PedsCom[32] consortia, *H. hepaticus*, SFB, *P. goldsteinii*, *L. monocytogenes*, *C. rodentium*, *S. Typhymurium*. Extracts were prepared either from agar plates or from liquid culture. For plate-grown bacteria, colonies were scraped from the agar with a sterile PBS-moistened swab into 5 mL sterile PBS, pelleted at 10,000 x g for 10 min, and resuspended in 1 mL sterile PBS. For liquid culture, 1.5 mL of culture grown from single colonies was pelleted at 10,000 x g for 10 min at 4 °C and resuspended in 200-500 µL sterile 1x PBS. In both cases, $OD_{600}$ was measured, and the suspensions were normalized to $OD_{600}$ ~1, so that all strains were used at a comparable concentration. To order bacteria by their approximate age of first colonization (Fig. 6C), two separate workflows using Claude Code (Anthropic) were used. The first assigned 34 hCom2 and PedsCom strains of our provisional colonization chart to one of four stages (pioneer/neonatal, early-infant/milk, transitional/weaning or late/adult-associated) or flagged as uncertain. Assignments came from a structured literature review run as a multi-agent workflow in Claude Code (Anthropic): for each organism, one agent compiled a dossier from PubMed (NCBI E-utilities) and web searches; a second agent independently re-checked the timing evidence, the host (mouse pup or human infant), any extrapolation from genus to species, and every cited reference (PMID, title, journal and year); and a final synthesis reconciled the two and enforced consistency within each genus. The review moved 12 of the 34 organisms to a different stage and graded the evidence as high (7), medium (23) or low (4) confidence. The second workflow was modeled similarly but using Claude Science (Anthropic) with new agents and additional bacteria that were not included in the first analysis. Species-level timing data are sparse, so many assignments rest on genus- or guild-level succession in human infant cohorts; the six PedsCom strains were staged from the pre-weaning mouse community they form, taken as analogous to human early life. Oligo-MM12 strains and the enteric pathogens and pathobionts were not part of this classification and are grouped separately. The classification, confidence grades, and supporting references (Table S6) were checked manually against the primary literature. For reactivity assays, bacterial strains were added at 1 µL per well in a final assay volume of 200 µL.

***In vitro* reactivity screening of T cell hybridomas**

Five to ten days prior to the NFAT-GFP reporter assay, CD45.1 C57BL/6 mice were injected subcutaneously with $1x10^6$ B16-Flt3L melanoma cells[54] to enrich for dendritic cells (DCs) in the spleen. Spleens from tumor-injected mice were harvested no more than 3 weeks post-injections. On the day of the experiment, spleens were digested in 400 U/mL collagenase D. Digested tissue was smashed against 70µm strainer and single-cell suspensions were treated with Red Blood Cell Lysis Buffer. $CD11c^+$ dendritic cells were then enriched using mouse CD11c MicroBeads UltraPure (Miltenyi Biotec). $5x10^4$ splenic dendritic cells (DC) were cultured in 100 µl in a 96-well U-bottom plate and pulsed with antigen extracts (luminal contents: 1 µL/well of 200 µL stock; bacterial extracts: 1 µL/well of $OD_{600}$ ~1, food: 50 µg/well) of interest for 2-4 h at 37°C in a 5% $CO_2$ incubator. Next, $1x10^4$ TCR-transduced NFAT-GFP cells were co-cultured in a 96-well plate with the pre-pulsed DCs and incubated at 37 °C in a 5% $CO_2$ for 40h. After the co-culture, cells were stained for live–dead zombie NIR, TCRβ-PE and CD4-APC. We evaluated TCR stimulation by measuring GFP expression in viable $TCR\beta^+$ $CD4^+$ hybridomas. MHC-II dependence for TCR reactivity was confirmed by incubating DCs with anti-mouse MHC-II (I-A/I-E) antibody (25 µg/mL, Bio X Cell). Strains that resulted in GFP signal above background (no Ag stimulation) in both the presence and absence of anti-MHC-II blocking were excluded from the assay. TCR responsiveness of each NFAT-GFP hybridoma line was determined by anti-CD3 stimulation (0.5 μg/well).

*Data Analysis of reactivity screening of T cell hybridomas*

TCR reactivity was quantified by GFP expression in viable TCRβ$^{+}$ CD4$^{+}$ hybridomas. Each clone was scored by $\Delta = \%GFP^{+}_{antigen} - \% GFP^{+}_{no\ antigen}$. The threshold of positive reactivity was based on the maximum Δ %GFP$^{+}$ experienced by our negative control TCR (mispaired TCRα and β). The maximum Δ %GFP$^{+}$ the control experienced across all screens was 4.52. Therefore, the threshold of positive reactivity was a Δ %GFP$^{+}$ of 5. When an antigen was tested in more than one experiment, Δ %GFP$^{+}$ was calculated against each experiment's own controls, and then the values were averaged.

**Cell sorting for scRNA-seq of the whole neonatal mouse small intestine**

To generate the single cell RNA-sequencing (10X Genomics) dataset on the neonatal murine small intestine, bulk intestinal tissue was profiled, by combining sorted subsets from three isolation protocols (ISCs, LECs, T cells) that on the day of the experiment were performed in parallel:

*Intestinal stem cell (ISC) isolation*

To enrich for small intestinal ISCs, the last ⅔ of the small intestine of each mouse ($n$ = 8 P6-P7 *Lgr5-EGFP-IRES-CreERT2* neonates) were used to isolate crypts as described above (Small intestinal epithelial cell sorting for rt-qPCR). All fractions were spun down at 400 g for 5 minutes, resuspended in FACS buffer (PBS + 2% FBS) with TruStain FxC block (1:1000), filtered and stained for 20 minutes at 4°C in the dark with the following antibodies: CD45-APC-Cy7, CD31-PE-Cy7, CD326-APC, PDPN-PerCp-Cy5.5 and CD24-PE (all at 1:200) in addition to the Lgr5-EGFP reporter signal. Samples were washed twice in FACS buffer and resuspended at 5-6 X10$^{6}$ cells/ml in FACS buffer with DAPI (33 ng/ml). For single-cell RNA-sequencing of the small intestine 20,000 LGR5$^{+}$ ISCs (alive, CD45$^{–}$, CD31$^{–}$, Lgr5-EGFP$^{+}$ singlets) and 5000 CD326$^{+}$ epithelial cells from this isolation were spiked into a mix of 25 000 bulk intestinal cells, 25,000 lymphatic endothelial cells and 25,000 CD3$^{+}$ T cells from the lymphatic endothelial cell and T cell isolations of the neonatal small intestine that were done in parallel (see below). Cells of this mix were resuspended at a final concentration of 1200 cells/μl.

*Intestinal lymphatic endothelial cell (LEC) isolation*

For bulk intestinal and lymphatic endothelial cell sorting experiments postnatal C57BL6/J mice, age-matched with mice from ISC sorting experiments, were utilized. The entire small intestine per mouse ($n$ = 20 C57BL6/J mice, males and females) was dissected, washed and cut into ~5 mm fragments into a 50 ml conical tube filled with PBS for additional washes. Tissue pieces were transferred into 25 ml of wash solution (RPMI (Gibco) with 2% FBS, 100 μg/ml Penicillin-Streptomycin, 25 mM HEPES buffer and 2 mM L-Glutamine) with 1 mM EDTA pH 8.0 and 1 mM DTT for shaking at 225 rpm and 37°C for 15 minutes. The supernatant after shaking was discarded and tissue fragments were resuspended in 8 ml advanced DMEM/F12 media with 100 μg/ml Penicillin-Streptomycin, 25 mM HEPES and 2 mM L-glutamine as well as DNase I (0.1 mg/ml) (Roche), 0.25% collagenase (Sigma) and 2 U/ml Dispase (Gibco). Two ceramic beads were added per tube to mince the tissue during a 10-minute incubation at 37°C and 225 rpm. The solution was filtered through a 70 μm cell strainer, spun at 1300 rpm and the cell pellet was resuspended in FACS buffer. Cells were stained for 20 minutes at 4°C in the dark with the following antibodies: CD45-APC-Cy7, CD31-FITC, PDPN-PE, LYVE1-ef660 and CD326-PE-Cy7 (all at 1:200). Cells were washed twice and resuspended at a final concentration of 5-6 million cells/ml for sorting. For scRNA-seq of the small intestine 25 000 lymphatic endothelial cells (alive $CD45^{-}$, $CD31^{+}$, $PDPN^{+}$, $LYVE1^{+}$ singlets) were spiked into 25 000 bulk intestinal cells from the same sample prep and mixed with the resulting sorted populations of the other isolations, as mentioned above. Cells were resuspended at a final concentration of 1200 cells/μl.

*Total T cell isolation*
We used our immune isolation protocol for neonates, as outlined earlier (n=8 C57BL/6 mice). Following PerColl-based enrichment of lamina propria lymphocytes, the cells were resuspended in FACS buffer with TruStain FcX block (1:1000) and stained with the following antibodies: CD45-APC-Cy7 (1:200), CD90.2-AF488 (1:500), CD3e-BV421 (1:100), TCRβ-AF700 (1:200) and CD4-PE-Cy7 (1:400). Instead of DAPI, 7-AAD (1:200, Biolegend) was used to gate out dead cells. 25 000 CD3+ T cells (alive $CD45^{+}$ $CD90^{+}$ $CD3^{+}$) were spiked into the cell suspension mix from the additional isolations above. The mix of cells was spun down at 1500 rpm, resuspended in 100 μl PBS + 2% fetal bovine serum (FBS) and submitted for 3' gene expression library preparation and sequencing (Nextseq 2000, P2 1x100).

**Single cell RNA-sequencing (10X Genomics) of timestamped small intestinal $CD4^{+}$ T cells**

*Single-cell RNA-seq library construction and processing*
$Tomato^{+}$ $CD4^{+}$ T cells were sorted from the small intestine IE, LP compartments and from mesenteric lymph nodes (mLN) (as described in "staining and gating strategies" above). Samples were hash-tagged by mouse, tissue, and age with TotalSeq-C anti-mouse hashtag antibodies (BioLegend). Gene expression, V(D)J T-cell receptor and cell-hashing libraries were prepared with the Chromium Next GEM Single Cell 5′ v2 workflow by The Rockefeller University Genomics Resource Center. They were sequenced on Illumina NextSeq 2000 (P3 flow cells, 2 lanes each; 26 cycles for read 1, 10 + 10 for the indices and 92 for read 2). For the March 2024 capture, the gene-expression matrix used in this study was generated from lane 1 only (the hashtag library from both lanes). The hashtag assigned to each mouse, tissue and age in each capture is listed in Table S1.

*Data processing of single-cell RNA-seq and TCR-seq libraries*
Raw fastq files were processed with cellranger count (v7.0.1) using the --libraries and --feature-ref options, so that gene expression and hashtag (Antibody Capture) reads were quantified together. Hashtag barcodes and fastq files are available at GEOXXXXXX. Reads were aligned to a custom mouse reference (GRCm39/mm39 with the tdTomato sequence added, built with cellranger mkref). Chemistry was detected as single cell 5′ R2-only, and intronic reads were counted. TCR libraries were assembled and annotated with cellranger vdj (v7.0.1) against the 10x Genomics GRCm38 V(D)J reference (v7.0.0). The five TCR outputs were combined with cellranger aggr (v7.0.1), which assigns clonotype identifiers across all captures.

*Quality control, demultiplexing and clustering*
Filtered feature-barcode matrices were analyzed in R with Seurat (v5). Hashtag counts were normalized by centered log-ratio (CLR), and cells were demultiplexed with HTODemux (positive quantile of 0.99). Only singlets were kept; cells with more than one hashtag (doublets) or with no

hashtag above background (negatives) were removed. Mitochondrial content was low (median 1.6% of reads, fewer than 0.1% of cells above 10%) and was regressed out when each capture was normalized with SCTransform. The first 20 principal components were used for UMAP and for Louvain clustering at resolutions from 0.1 to 1. The five captures were merged, using variable genes selected across captures with SelectIntegrationFeatures, and clustered. Three clusters of the merged object (resolution 0.3) with low TCR recovery were removed: two expressed B-cell or epithelial genes (12% and 3% of cells with a TCR) and one consisted of T cells with low TCR recovery (42%), whereas at least 80% of cells carried a TCR in every other cluster. The remaining 34,483 αβ T cells were re-normalized with SCTransform and re-clustered as above, and the final resolution (0.2, nine clusters) was chosen for biological separation of T-cell states. Clusters were annotated by enrichment analysis (clusterProfiler v4.6.2, fgsea v1.24.0) and manual curation as TFH, Treg, naïve, cytotoxic/effector, adult IEL, IFN-induced (LN), TRM precursors, TFH/tissue remodeling and proliferating cells. Module scores for curated gene sets (AddModuleScore in Seurat; gene sets are listed in Table S3 and in the code repository) were compared across clusters with the Kruskal-Wallis test, followed by pairwise Wilcoxon tests with Benjamini-Hochberg correction. Differential expression between weaning cells and suckling or adult cells was tested separately for LP and IE with the FindMarkers function in Seurat (Wilcoxon rank sum test), and genes with Bonferroni-adjusted p-values smaller than 0.05 were considered significant. Full settings are listed in the code repository. For single-cell heatmaps, expression of each gene was z-scored across the cells shown, and cells were ordered by age and cluster (Fig. 3F) or by age group and clone (Fig. 6D)

*TCR repertoire analysis*
Plate-based TCRβ sequencing was performed as described in Bilate et al.[38] Fasta files were aligned and analyzed on IMGT[55] (https://www.imgt.org/HighV-QUEST/). For each mouse and tissue with at least 50 retained cells, clones were ranked by frequency (% of cells). Frequencies were averaged across mice at each rank (mean ± SEM) and shown only where at least two mice contributed. Mean curves were made non-increasing by a running minimum over rank, which removes increases caused by mice with fewer clones dropping out at deeper ranks. Diversity 50 (D50) was calculated by ranking clones by frequency, as the fraction of clones needed to account for 50% of the cells of each mouse and tissue. Repertoire overlap between tissues of the same mouse was quantified with the Morisita-Horn index computed on clone counts (plug-in estimator for the 10X data, as implemented in the code repository; divo R package for the plate-based data). Circos plots were created with Circos (v0.69-8). For each mouse, each tissue is an arc whose length is its number of cells, each clone is a band whose width is its number of cells, and ribbons connect clones shared between tissues. Graphs were generated in R (v4.5.0) with Seurat (v5.3.1), ggplot2 (v4.0.2), ggrastr (v1.0.2), ggrepel (v0.9.8) and ComplexHeatmap (v2.25.0). Figures were assembled in Adobe Illustrator (RRID:SCR_010279). Statistical tests are indicated in the figure legends, and p values smaller than 0.05 were considered significant.

**Single cell RNA-sequencing (10X Genomics) of the neonatal (P6-P7) small intestine**

FASTQ files were generated by the sequencing core with Illumina bcl2fastq. Cell Ranger count (v6.0.1; chemistry detected as Single Cell 3′ v3; intronic reads not counted) was used to align reads to the GRCm38 (mm10; 10x Genomics mm10-2020-A) reference genome with the EGFP-IRES-CreERT2 sequence added and to generate count matrices. The adult sample (Niec et al.[19];

GSM5712427) had been processed with the same Cell Ranger version and reference, and its raw count matrix was used here.

*Preprocessing and integration of single cell RNA-sequencing datasets.*
We first used CellBender[56] to remove ambient RNA molecules from the raw Cell Ranger count matrices (remove-background, --fpr 0.01, --epochs 150; P6: CellBender v0.3.0, --expected-cells 12000; adult: an earlier CellBender release run in 2022, --expected-cells 5000 --total-droplets-included 20000). All other parameters were left at their defaults. The resulting filtered cellbender.h5 file was then loaded into R (v 4.4.1) and single cell data were analyzed with Seurat (v5.4.0) following the Seurat v5 vignette[57]. Seurat objects were created for the postnatal and adult small intestinal dataset (as published by Niec et al.[19]) and processed in parallel. Quality metrics included subsetting objects for nFeature_RNA > 200 and < 8000 and a mitochondrial percentage of less than 15%. The data were normalized (LogNormalize) followed by variable feature detection ("vst", nfeatures = 3000). Doublets were excluded following doublet scoring via scrublet[58] (loaded in a conda environment via reticulate) before the two datasets were merged. In the merged object the counts were split into one layer per dataset, re-normalized, 3,000 variable features were selected, and the data were scaled and analyzed by PCA with default settings. Subsequently the data layers were integrated using the RPCA integration, as recommended by the Seurat team (following the general workflow, as described here: https://satijalab.org/seurat/articles/seurat5_integration). After that, we ran FindNeighbors (dims = 1:20), FindClusters (resolution = 2; 40 clusters) and RunUMAP (dims = 1:20) on the RPCA-integrated reduction. To identify the top marker genes per cluster and label cell types, we performed differential gene expression with the FindAllMarkers function (only.pos = TRUE, min.pct = 0.25, logfc.threshold = 0.25). The 40 clusters were annotated, based on top marker genes and manual curation, as 23 cell types (9,048 P6 and 2,636 adult cells; proliferating ISCs are labelled TA cells in Fig. 1G,H), renamed and cell types were ordered roughly according to their location along the crypt-villus axis in the tissue for the dot plot in Fig. 1H. Plots were visually enhanced by using ggplot2[59] and the scCustomize (v3.2.4)[60] package in R, unless specified otherwise.

***Differential gene expression analyses***
To perform differential gene expression analysis on overlapping cell type across postnatal timepoints in our integrated dataset, we pooled the subclusters of the respective cell type (e.g., epithelial cells) and ran FindMarkers on the log-normalized RNA assay (Wilcoxon rank sum test; logfc.threshold = 0.25, min.pct = 0.1) for the two groups of interest. Genes with Bonferroni-adjusted $p < 0.05$ and $|\log2 \text{ fold change}| \geq 0.585$ were considered significant, and genes whose symbols contain "Rp" (mostly ribosomal protein genes) were removed. Quiescent ISCs, TA cells (proliferating ISCs), secretory progenitors and Paneth cells (adult only) were classified as crypt epithelial cells, and enterocytes, goblet cells and enteroendocrine cells as villus epithelial cells; tuft cells (adult only) were not included in either group. For Suppl. Fig. 1G, all epithelial cell types were pooled (including the single P6 Paneth cell and the adult tuft cells). For Fig. 1I, the average log2 fold change (adult vs. P6) of nine genes from the Fig. 1H panel (H2-Ab1, H2-DMb1, H2-Aa, Cd74, Prdm1, Ccl25, Cxcl16, Il15, Il18) was plotted in GraphPad Prism for villus and crypt epithelial cells in which the difference was significant (adjusted $p < 0.05$ and $|\log2 \text{ fold change}| \geq 0.585$); the villus Cxcl16 bar is shown although it was not significant (adjusted $p = 1.0$). The

volcano plot of DEGs was built by inputting our list of DEGs into Claude. Data visualization was independently validated by plotting in R, using ggplot2[59].

***Integration of single cell RNA-sequencing with published spatial transcriptomic (10X Visium HD)***

To project resolved timestamped $CD4^+$ T cell clusters (as indicated in figure legend) onto spatial transcriptomic data of the murine adult small intestine, we used 10X Genomics' publicly available FFPE 10X Visium HD data set on the mouse small intestine at steady state ((link: https://www.10xgenomics.com/datasets/visium-hd-cytassist-gene-expression-libraries-of-mouse-intestine) (SI of 8wko C57BL/6 mice (FFPE, Visium CytAssist, mouse transcriptome probe set v2.0 (43x50)), 10x Visium HD ST dataset by space ranger v3.0, 10X Genomics (2024/3/25)). In short, the filtered_feature_bc_matrix.h5 file of 10X Visium HD data was imported into R (v 4.4.1) and processed via Seurat according to default settings. We performed our analyses on the 8 µm bin data, normalized (LogNormalize),ran FindVariableFeatures, scaled the data and performed PCA, FindNeighbors (dims = 1:30), FindClusters and UMAP (dims = 1:30) analyses as described for single cell studies above. To generate a matching and comprehensive single cell reference, we had originally integrated our joint postnatal/adult dataset from above with our timestamped T cell data (only IE and LP data) as well as an additional enterocyte dataset, deposited at GSM8677456 and GSM8677457[61]. The two enterocyte bioreplicates were processed as described: We first used CellBender[56] to remove ambient RNA molecules from the raw Cell Ranger count matrices (remove-background, --fpr 0.01, --epochs 150; P6: CellBender v0.3.0, --expected-cells 500). All other parameters were left at their defaults. Those enterocyte data were then processed in Seurat as described for postnatal/adult whole intestinal data. Similarly, the already annotated fullmeta.rds file of our timestamped T cell (10X Genomics) single cell data were imported into R and re-processed. We subset the data by excluding mLN data, LogNormalized (based on the RNA assay) and ran FindVariableFeatures (selection.method = "vst", nfeatures = 3000). We double-checked for and excluded doublets following doublet scoring via scrublet[58] (loaded in a conda environment via reticulate). The enterocyte dataset, joint postnatal/adult whole intestine and subset timestamped T cell datasets were then merged as described above. In the merged object the counts were split into one layer per dataset, re-normalized, 2,000 variable features were selected, and the data were scaled and analyzed by PCA with default settings. Subsequently the data layers were integrated using the RPCA integration, as recommended by the Seurat team. After that, we ran FindNeighbors (dims = 1:30), FindClusters (resolution = 2; 45 clusters) and RunUMAP (dims = 1:30) on the RPCA-integrated reduction. To identify the top marker genes per cluster and label cell types, we performed differential gene expression with the FindAllMarkers function (only.pos = TRUE, min.pct = 0.25, logfc.threshold = 0.25). The 45 clusters were annotated, based on top marker genes and manual curation, as 30 cell types. To generate a reference for the spatial Visium HD data, we subset that integrated dataset on our timestamped T cell clusters (IE and LP data pooled per timepoint) and merged 3-7 and 6-10 wko into one common adult cluster, resulting in three clusters: P1-P16 (1250 cells), P1-P22 (4220 cells) and Adult (2950 cells). We subset our whole integrated dataset to those T cell clusters, collapsed per timepoint, since otherwise more abundant cell populations would outpower T cells when projecting single cell onto spatial data. However, we intentionally kept the original PCA reduction to better distinguish T cell bins in the spatial data matrix from non-immune/T cell bins, which would not have been possible by simply using our timestamped T cell data alone (not embedded in a whole-tissue dataset) as our reference. To integrate single cell with spatial date, we employed Seurat's label transfer workflow and ran

FindTransferAnchors, specifying our subset single cell object as our reference and the processed Visium HD input data as our query object (reference.reduction = "pca", dims = 1:30). Using the resulting anchors as our anchorset, we employed TransferData (dims = 1:30, k.weight = 32) and added the resulting predictions of bins containing any of the indicated 3 T cell clusters to the metadata of our Visium HD object. We extracted and thresholded the prediction scores (> 0) per cluster to only plot the top 10% of bins predicted to contain any of the T cells from our single cell reference clusters (in a "yes" or "no" fashion). These newly assigned categories were subsequently plotted via the SpatialDimPlot function. We confirmed overlapping patterns when plotting the entire set of prediction scores (gradient) on spatial feature plots. In sum, we projected selected cell types from our single cell dataset onto 10X Visium HD data to infer where T cells of various stages would most likely localize in the tissue, with the caveat that spatial data were merely available from the adult small intestine and thus not matched by timepoints. We focused on T cell clusters of interest to better resolve T cells located in spatial bins, while the abundance of epithelial and stromal cells in our original single cell reference would have likely skewed projections towards those cell types had we input our entire single cell dataset as our reference.

*SpaceFold projection of deconvolved 10X Visium spatial transcriptomic data.*
To generate spatial gene expression plots in supplementary figure 4D (*Gzmb* and *Foxp3*) the integrated transcriptomic data from Niec *et al.* (2022) were used and re-analyzed[19]. Briefly, 10X Visium spatial transcriptomic data on the adult mouse small intestine were deconvolved using the matched adult SI single cell RNA-sequencing reference, which we also used in this paper. We used the published deconvolution of these data with BayesPrism[62], available on figshare, followed by dimensional projection via SpaceFold as previously described[19]. Expression of *Gzmb* and *Foxp3* was only plotted on T cells, which included total T cells as recovered from intestinal tissue but without any sort of prior enrichment, and was normalized (as part of the algorithm) on a per cell basis.

**Data and code availability**
Raw and processed sequencing data, the per-capture matrices and the annotated Seurat object are available at GEO. All code used to generate analyses and figures are available upon request and will be available at github.com/tiagobrc/Bilate2026.

**Quantification and statistical analysis**
Data are expressed as individual values per mouse and mean ± standard error of the mean (SEM) and were pooled from at least 2 independent experiments as specified in figure legends. The number of biological and/or technical replicates per experiment is indicated in each figure legend. For immunofluorescence studies at least three images per biological replicate were taken, analyzed and averaged (except for T cell z-position analyses as specified). Image analyses were done in Imaris v.9.5-10 (Bitplane) unless specified otherwise in the respective Methods section. Flow cytometry analyses were done using FlowJo™ v.10 software. Statistical analyses were performed using GraphPad Prism v.11 software. Only conditions of interest were compared, as indicated in the figure legends. Unpaired two-tailed Student's t-tests were used for comparisons between two groups. Paired two-tailed Student's t-tests were used for selected data from *Sell-Cre*$^{ERT2}$ *ROSA26*$^{LSL\text{-}tdTomato}$ mice, where Tomato$^{+}$ and Tomato$^{-}$ cells were matched per animal. One-way analysis of variance (ANOVA) with Tukey's correction for multiple comparisons was used to

assess statistical significance between three or more groups with one experimental parameter. Graphs were plotted using Prism v.11 or R and edited for appearance using Adobe Illustrator 2025.

**AI-assisted analysis and practices**

Generative AI was used as a coding and research assistant under the authors' supervision. Claude Code (Anthropic) was used to write and refactor the R and Python analysis and plotting code for the single-cell and TCR repertoire figures (module scores, heatmaps, volcano plots, clonotype rank-abundance curves and Circos plots); to assemble and document the code repository; to re-run the processing pipelines from the raw sequencing data as a reproducibility check; to cross-check these methods against the code and data; and to run the literature review of bacterial colonization timing described above. Claude Science (Anthropic) also facilitated the literature review of bacterial colonization timing in addition to the analysis and generation of heat maps in Figure 6. Claude (Anthropic) was also used to shortlist candidate chemokine, MHC-II and interleukin genes from the differential-expression lists for Fig. 1H, to assign differentially-expressed genes into gene categories, and subsequently prepare the volcano plot in Suppl. Fig. 1G. AI tools did not generate or alter data. The panels of Fig. 1G-I, Fig. 3, Fig. 5C, 5F and 5G, Fig. 6D and Suppl. Figs. 1E-G, 3B-D, 5A-C and 5F are produced by deterministic scripts (github.com/tiagobrc/Bilate2026) from the processed objects deposited at GEO; these scripts reproduce the original analysis outputs, and the rank-abundance statistics were also re-derived in an independent implementation. Flow cytometry, imaging, and the NFAT reactivity screen were analyzed as described in the corresponding sections. The authors reviewed all AI-generated code, text, and literature summaries and take full responsibility for the content.

### SUPPLEMENTARY FIGURES

#### Figure S1

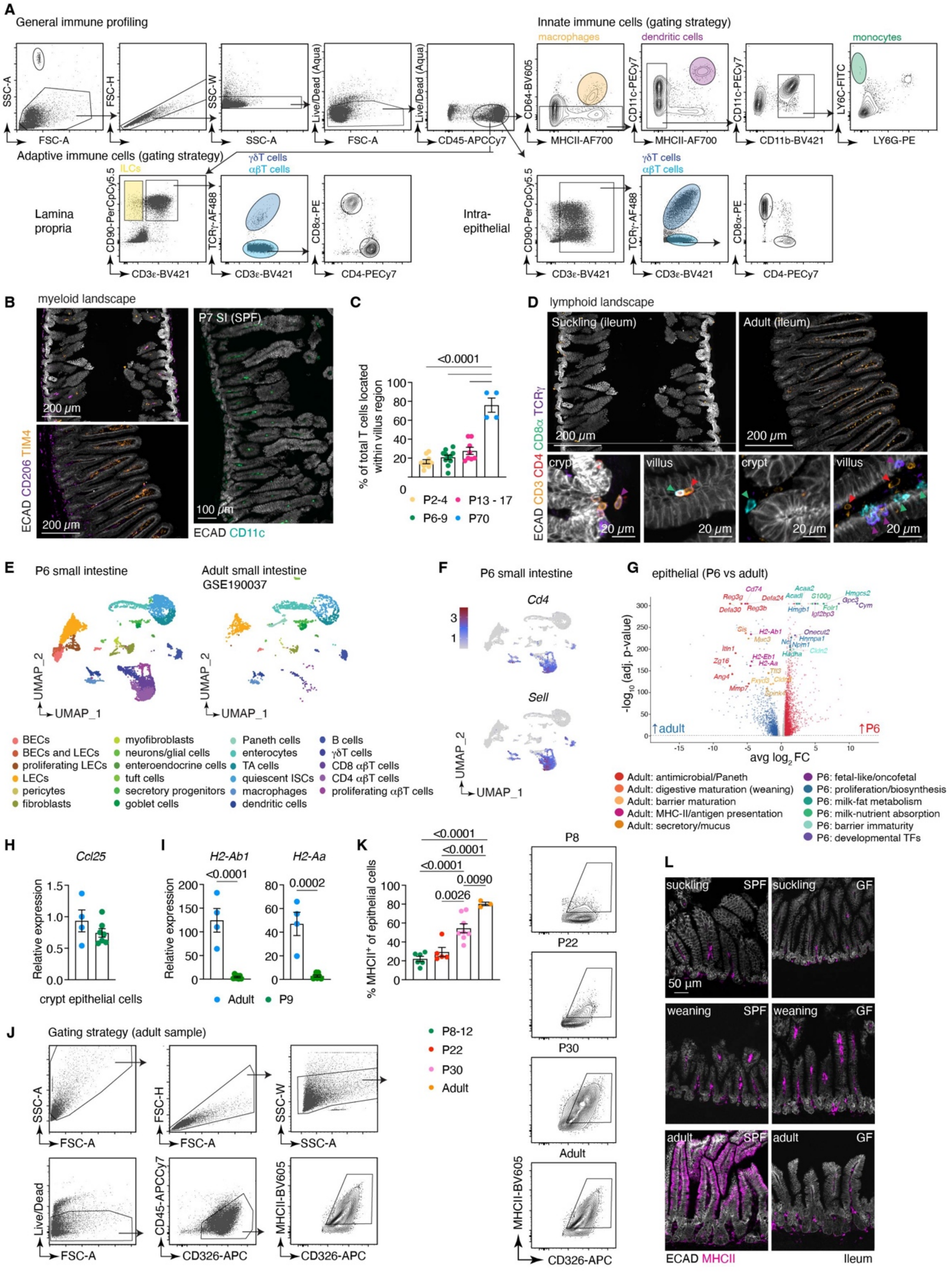

**Figure S1. (related to Figure 1)**

**(A)** Gating strategy for general immune profiling shown in figure 1A. **(B-D)** Multiplex immunofluorescence staining for myeloid (B) and lymphoid (D) immune subsets in the ileum of suckling (P8) and adult SPF mice. Crypt and villus zoom-ins are below whole-tissue images in D. **(C)** Quantification of T cells in the villus of SPF mice of whole-mount images shown in figure 1C. **(E)** Separate UMAPs visualizing mouse small intestinal scRNA-seq data from postnatal (P6-7, left) and adult (right, re-analyzed from GSE190037) samples. **(F)** Feature plots (*Cd4*, *Sell*) in postnatal small intestinal data. **(G)** Volcano plot showing differentially expressed genes (DEGs) in P6 versus adult epithelial cells from scRNA-seq data in E. Genes are color-coded by the categories listed below. **(H-I)** Relative expression of *Ccl25* (H), *H2-Ab1* and *H2-Aa* (I) via rt-qPCR on small intestinal crypt epithelial cells of neonates (P9) and adults. One dot per mouse. n = 4-7 mice/age. **(J)** FACS gating strategy to quantify MHC-II$^+$ epithelial cells, as shown in K. **(K)** Representative flow plots of MHC-II$^+$ CD326$^+$ small intestinal epithelial cells across postnatal timepoints. The percentage of MHC-II$^+$ epithelial cells is shown in the graph on the top left (1 dot per mouse; n = 3-8 mice/age). **(L)** Representative sagittal sections illustrating epithelial and non-epithelial MHC-II expression in the ileum of suckling (P7), weaning (P18) and adult (8 weeks of age) SPF and GF animals. Each symbol represents a mouse (C, H, I, K). One-way ANOVA with Tukey's multiple comparisons-test was used for data in panels C and K, an unpaired two-tailed Student's t-test was used to compare groups in H-I. Where applicable, graphs show mean ± the standard error of the mean (SEM).

**Supplementary video 1 (related to figure 1).** (video Exp9_CD3_Ecad_Lyve1_dSI_P9-1_003)

Animated view of the small intestinal (ileum) crypt-villus axis from bottom (below the crypts, as indicated by the lymphatic vasculature (LYVE-1$^+$, green)) to top. The underlying sample is a cleared whole-mount ileal sample from a neonatal (P9) C57B6 wild-type mouse. Epithelial cells (ECAD$^+$, red) and CD3$^+$ T cells (white) are shown. The video is representative of *n* = 5 mice.

**Supplementary video 2 (related to figure 1).** (video Exp9_CD3_Ecad_Lyve1_adult_1_002)

Animated view of the small intestinal (ileum) crypt-villus axis from bottom (below the crypts, as indicated by the lymphatic vasculature (LYVE-1$^+$, green)) to top. The underlying sample is a cleared whole-mount ileal sample from an adult (10 weeks of age) C57B6 wild-type mouse. Epithelial cells (ECAD$^+$, red) and CD3$^+$ T cells (white) are shown. The video is representative of *n* = 4 mice.

#### Figure S2

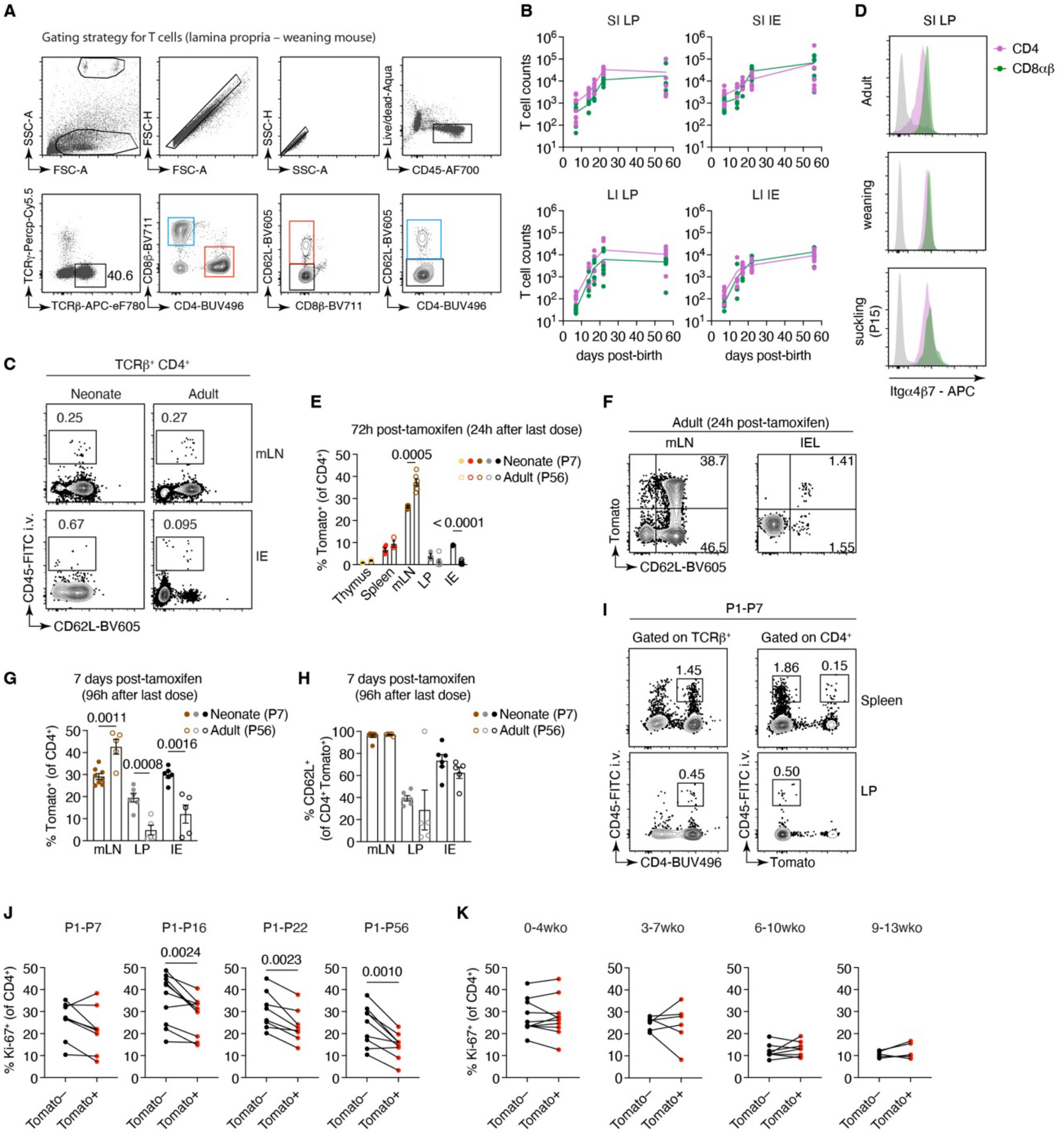
A
Gating strategy for T cells (lamina propria – weaning mouse)
SSC-A
FSC-A
FSC-H
SSC-H
Live/dead-Aqua
CD45-AF700
TCRγ-Percp-Cy5.5
TCRβ-APC-eF780
40.6
CD8β-BV711
CD4-BUV496
CD62L-BV605
CD8β-BV711
B
SI LP
SI IE
LI LP
LI IE
T cell counts
days post-birth
D
SI LP
CD4
CD8αβ
Adult
weaning
suckling (P15)
Itgα4β7 - APC
C
TCRβ+ CD4+
Neonate
Adult
0.25
0.27
mLN
0.67
0.095
IE
CD45-FITC i.v.
CD62L-BV605
E
72h post-tamoxifen (24h after last dose)
% Tomato+ (of CD4+)
0.0005
< 0.0001
Neonate (P7)
Adult (P56)
Thymus
Spleen
mLN
LP
IE
F
Adult (24h post-tamoxifen)
mLN
IEL
38.7
46.5
1.41
1.55
Tomato
CD62L-BV605
G
7 days post-tamoxifen (96h after last dose)
Neonate (P7)
Adult (P56)
0.0011
0.0008
0.0016
% Tomato+ (of CD4+)
mLN
LP
IE
H
7 days post-tamoxifen (96h after last dose)
Neonate (P7)
Adult (P56)
% CD62L+ (of CD4+ Tomato+)
mLN
LP
IE
I
P1-P7
Gated on TCRβ+
Gated on CD4+
1.45
1.86
0.15
Spleen
0.45
0.50
LP
CD45-FITC i.v.
CD4-BUV496
Tomato
J
P1-P7
P1-P16
P1-P22
P1-P56
0.0024
0.0023
0.0010
% Ki-67+ (of CD4+)
Tomato–
Tomato+
K
0-4wko
3-7wko
6-10wko
9-13wko
% Ki-67+ (of CD4+)

**Figure S2. (related to Figure 2)**

**(A)** Gating strategy for profiling T cell subsets shown in figure 2A-B and **(B)** absolute counts of lymphocytes from SI and LI LP or IE. (n=4-8 mice/age). **(C)** Representative flow cytometry plots of blood-derived naïve $CD4^+$ T cells in neonates and adults. **(D)** Histograms showing integrin $\alpha 4\beta 7$ among $CD4^+$ and $CD8^+$ $\alpha\beta$T cells in suckling (P15), weaning (P21) and adult (>P60) mice. Gray histogram, $CD45^-$ cells as reference. **(E)** Quantification of labelling of $CD4^+$ T cells across tissues in neonate (P7) and adult (P56) Sell-Tomato mice 72 hours post-tamoxifen. (n=3-6 mice/age) **(F)** Representative FACS plots of Sell-Tomato mLN or IE T cells 24h post-tamoxifen. **(G-H)** Quantification of Tomato labelling of $CD4^+$ T cells (G) and percentage of naïve timestamped $CD4^+$ T cells (H) 7 days post-tamoxifen. (n=4-8 mice/age). **(I)** Representative FACS plots showing the percentage of circulating $CD4^+$ and timestamped T cells in the spleen and LP of neonates. **(J-K)** Quantification of proliferation among timestamped and non-timestamped $CD4^+$ T cells across timestamping windows. Each symbol represents a mouse (E, G, H, J, K). Graphs show mean ± SEM. Statistical significance was determined by paired Student's t-test for J-K. (n=5-10 mice).

**Figure S3**

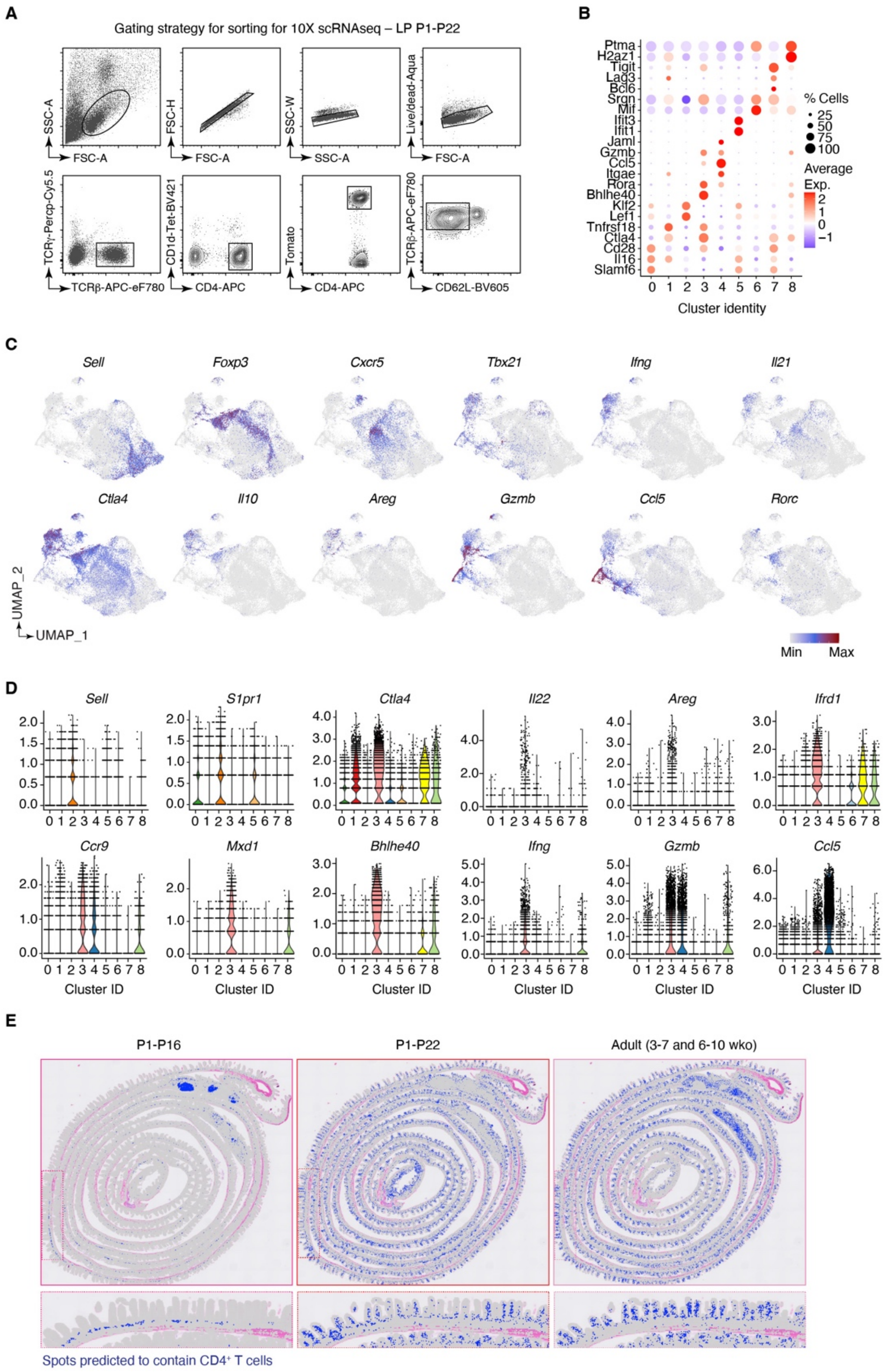
A
Gating strategy for sorting for 10X scRNAseq – LP P1-P22
SSC-A
FSC-A
FSC-H
SSC-W
Live/dead-Aqua
TCRγ-Percp-Cy5.5
TCRβ-APC-eF780
CD1d-Tet-BV421
CD4-APC
Tomato
CD62L-BV605
B
Ptma
H2az1
Tigit
Lag3
Bcl6
Srgn
Mif
Ifit3
Ifit1
Jaml
Gzmb
Ccl5
Itgae
Rora
Bhlhe40
Klf2
Lef1
Tnfrsf18
Ctla4
Cd28
Il16
Slamf6
% Cells
25
50
75
100
Average Exp.
2
1
0
−1
Cluster identity
C
Sell
Foxp3
Cxcr5
Tbx21
Ifng
Il21
Ctla4
Il10
Areg
Gzmb
Ccl5
Rorc
UMAP_2
UMAP_1
Min
Max
D
Sell
S1pr1
Ctla4
Il22
Areg
Ifrd1
Ccr9
Mxd1
Bhlhe40
Ifng
Gzmb
Ccl5
Cluster ID
E
P1-P16
P1-P22
Adult (3-7 and 6-10 wko)
Spots predicted to contain CD4+ T cells

**Figure S3. (related to Figure 3)**

**(A)** Gating strategy used for sorting of timestamped $CD4^+$ T cells for scRNA-seq**. (B)** Dot plot with top genes per cluster. **(C-D)** Expression of the indicated genes by timestamped T cells on UMAP (C) and per cluster (violin plot, D). **(E)** Spatial projection of time-stamped suckling (P1-16), weaning (P1-22) and adult (3-7 wko) $CD4^+$ T cells from scRNA-seq data onto 10X Genomics' publicly available Visium HD healthy mouse small intestine (8 μm spatial bins) dataset (see methods for details). Spots predicted to contain $CD4^+$ T cells (top 5%) are colored in dark blue. Top panels show the entire small intestine, while zoom-ins (of boxed regions) below illustrate predicted $CD4^+$ T cell spots along the crypt-villus axis.

Figure S4

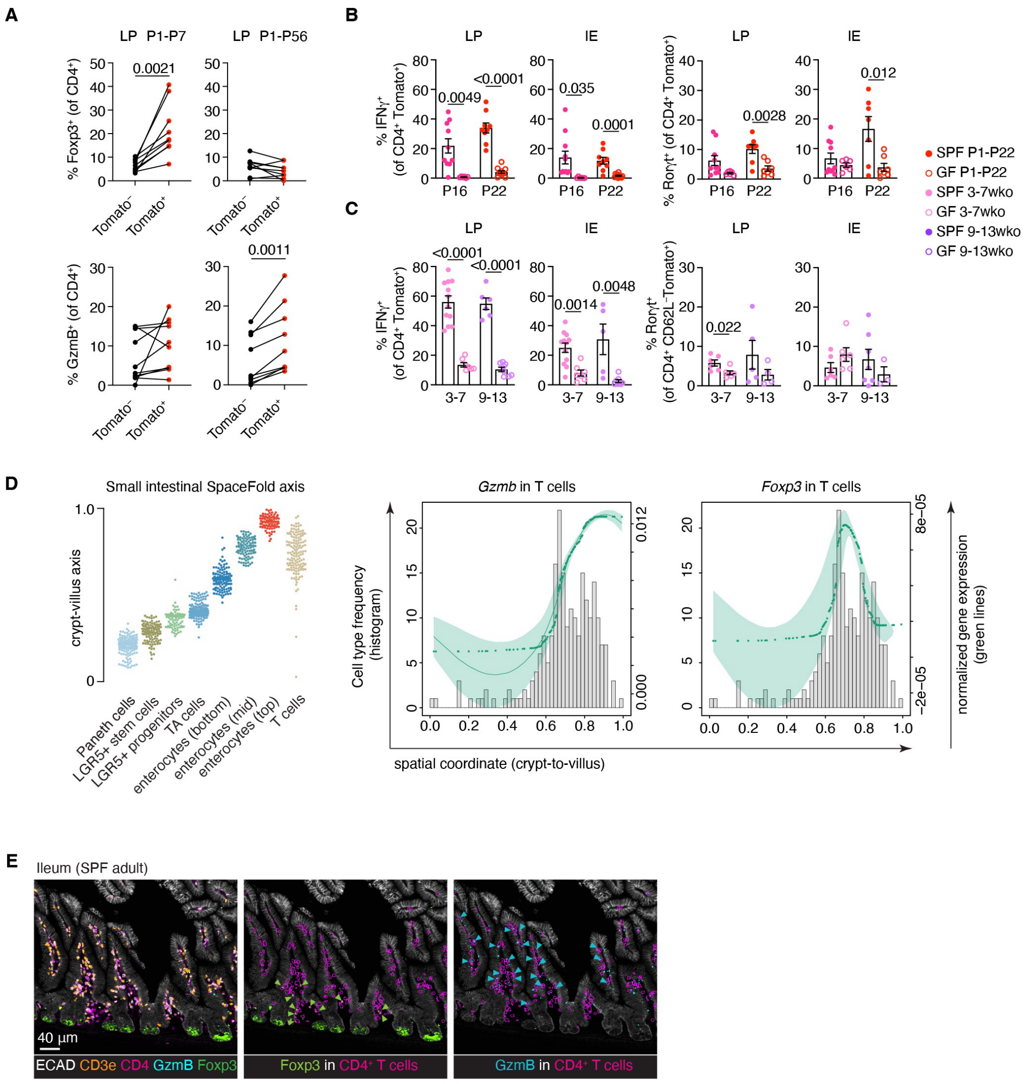

### Figure S4. (related to Figure 4)

**(A)** Foxp3 (top) and Gzmb (bottom) expression in timestamped and non-traced LP CD4$^+$ T cells in the indicated time windows, matched per animals. (n=9-10 mice) **(B-C)** Quantification of IFN$\gamma^+$ and Ror$\gamma$t$^+$ CD4$^+$ T cells in SPF and GF mice across timestamping windows. SPF mice depicted here are the same shown in Figure 1A**.** (n=3-12 mice/age). **(D)** Axis of ordered deconvolved cell types from adult small intestinal spatial transcriptomic data (left). Spatial gene expression plots of *Gzmb* and *Foxp3* in T cells along the small intestinal crypt-villus axis next to it (middle and right plots). The histogram indicates T cell frequency along the crypt-villus axis (from left to right), while the green line refers to normalized gene expression. **(E)** IBEX images of the ileum of adult (6-8 wko) SPF mice Left: Gzmb$^+$ and Foxp3$^+$ T cells in the ileum. Middle: Foxp3-expressing CD4$^+$

T cells. Green arrows indicate Foxp3$^{+}$ cells, while purple surfaces mark CD4$^{+}$ T cells. Right: Gzmb-expressing CD4$^{+}$ T cells. Blue arrows indicate Gzmb$^{+}$ cells, while purple surfaces mark CD4$^{+}$ T cells

Each symbol represents one mouse. Graphs in B-C show mean ± standard error of the mean (SEM). Statistical significance was assessed by paired (A) or unpaired (B-C) Student's t-test.

Figure S5

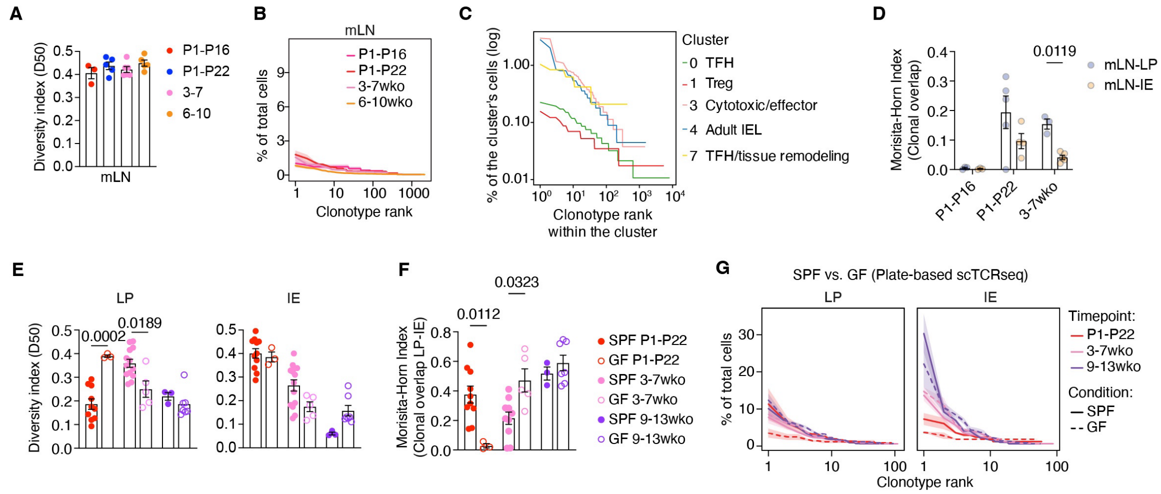


**Figure S5. (related to figure 5)**

**(A-B)** TCR diversity measured by diversity index 50 (A) and clonotype-ranking analysis (B) of CD4$^{+}$ T cell clones harvested from mLN across timestamping windows. (n=3-6 mice/age). **(C)** Clonotype-ranking analysis within individual clusters: 0 (TFH), 1 (Treg), 3 (cytotoxic/effector), 4 (adult IEL), 7 (TFH/tissue remodeling). **(D)** Clonal overlap between mLN and IE or LP determined by Morisita-horn index from mice at the indicated timestamping windows. (n=3-5 mice/age). **(E-G)** TCR Diversity (E), Morisita-horn (F) indices and clonotype-ranking analysis (G) of TCR clones in SPF and GF mice (n=3-13 mice/age). Each symbol represents a mouse (A, D-F). Graphs show mean ± SEM. Lines and shaded areas represent the mean and SEM of all mice included in the analysis. Statistical significance was determined by unpaired Student's t-test between matched groups. SPF mice shown in D-G are also shown in Figure 5 for comparison.

#### Figure S6

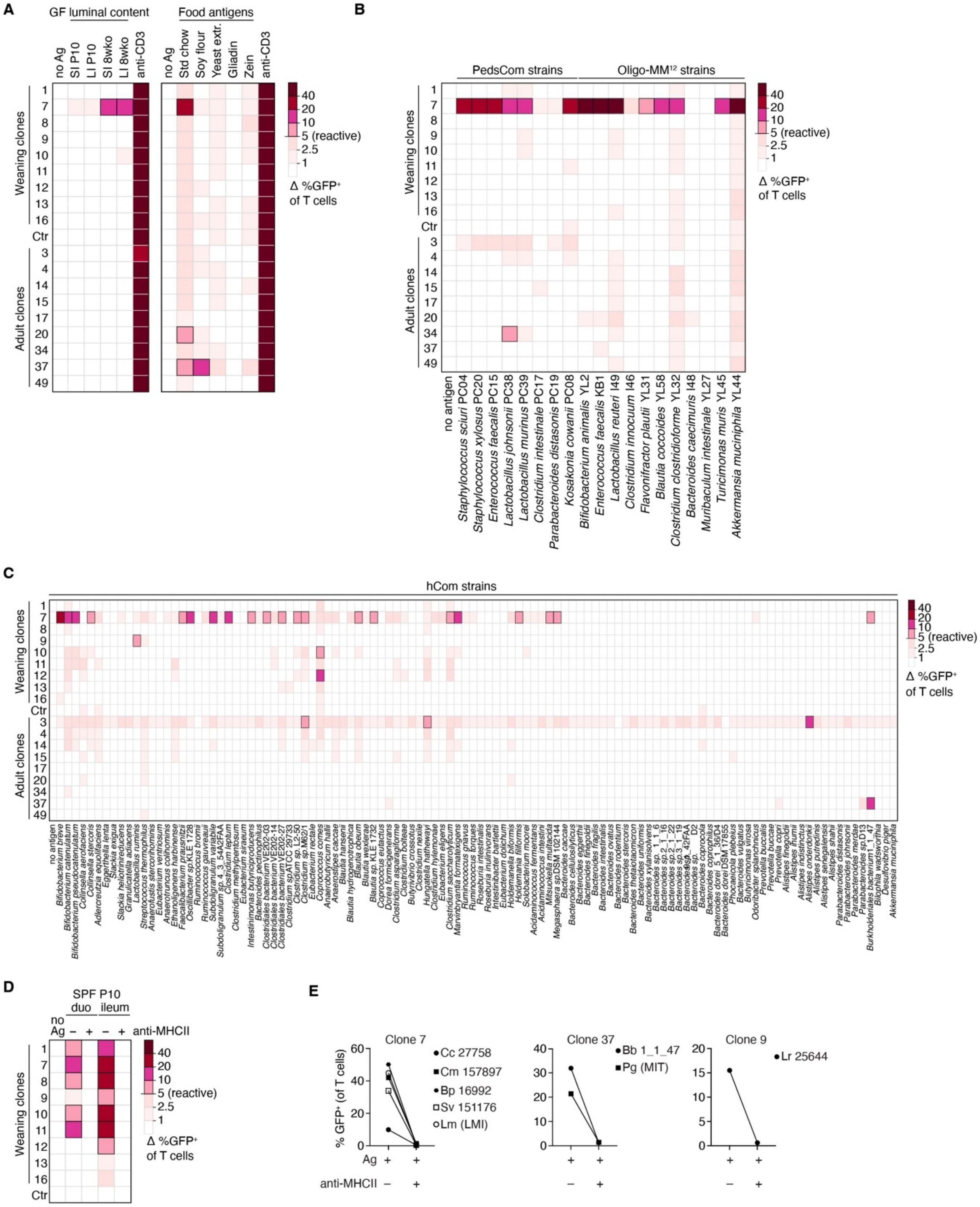

A
GF luminal content
Food antigens
no Ag
SI P10
LI P10
SI 8wko
LI 8wko
anti-CD3
no Ag
Std chow
Soy flour
Yeast extr.
Gliadin
Zein
anti-CD3
Weaning clones
Adult clones
Ctr
40
20
10
5 (reactive)
2.5
1
Δ %GFP⁺ of T cells
B
PedsCom strains
Oligo-MM12 strains
no antigen
Staphylococcus sciuri PC04
Staphylococcus xylosus PC20
Enterococcus faecalis PC15
Lactobacillus johnsonii PC38
Lactobacillus murinus PC39
Clostridium intestinale PC17
Parabacteroides distasonis PC19
Kosakonia cowanii PC08
Bifidobacterium animalis YL2
Enterococcus faecalis KB1
Lactobacillus reuteri I49
Clostridium innocuum I46
Flavonifractor plautii YL31
Blautia coccoides YL58
Clostridium clostridioforme YL32
Bacteroides caecimuris I48
Muribaculum intestinale YL27
Turicimonas muris YL45
Akkermansia muciniphila YL44
C
hCom strains
D
SPF P10
duo
ileum
no Ag
anti-MHCII
E
Clone 7
Cc 27758
Cm 157897
Bp 16992
Sv 151176
Lm (LMI)
Clone 37
Bb 1_1_47
Pg (MIT)
Clone 9
Lr 25644
% GFP⁺ (of T cells)
Ag
anti-MHCII

**Figure S6. (related to Figure 6)**

**(A)** Heatmaps show the reactivity of weaning and adult clones to luminal content from the small (SI) and large (LI) intestine of germ-free (GF) mice at P10 and 8 weeks of age (left), and to dietary antigens (right). No Ag, no antigen stimulation, Ctr, control TCR with mispaired TCRαβ, anti-CD3 stimulation was used as positive control for hybridoma performance. **(B)** Reactivity of the same clones to individual bacterial strains from the PedsCom and Oligo-$MM^{12}$ defined consortia. (**C)** Reactivity of clones to an extended panel of individual strains from the hCom2 defined human commensal community. No Antigen, no antigen stimulation. **(D)** Reactivity of weaning clones to duodenal and ileal luminal content from P10 SPF mice in the presence or absence of anti-MHC-II blocking antibody. Ctr, control TCR with mispaired TCRαβ. **(E)** Percentage of $GFP^+$ T cells of clones 7, 37, and 9 tested against their reactive strains in the presence or absence of anti-MHC-II blocking antibody.
